# Defining the Functional Consequence of the Human Ocular Microbiome

**DOI:** 10.64898/2026.10.28.762462

**Authors:** Jaishree Gandhi, Brinna Desai, Simran Mangwani-Mordani, Jackie Shane, Julia Bierker, Alysha Wessell, Daphne M. Rodriguez J., Simran Kaur Randhawa, Xinning Ma, Catherine M. Phelps, Marlies Meisel, Asha Patel, Kelvin Li, Adam Fitch, Barbara Methe, Vishal Jhanji, Anat Galor, Anthony J. St. Leger

**Author notes:** Authors contributed equally to this work (Jaishree Gandhi, Brinna Desai). Authors contributed equally to this work (Anat Galor, Anthony J. St. Leger). Correspondence: Anthony St. Leger Correspondence: Anat Galor. The authors have declared that no conflict of interest exists.

## Abstract

The ocular surface is one of the body’s most environmentally exposed mucosal barriers, yet it is considered low biomass and harbors orders of magnitude fewer bacteria than microbe-rich sites like the gut. This has cast doubt on whether a functional core ocular microbiome exists at all. Having previously defined a protective γδ T cell–IL-17 axis engaged by the murine commensal *Corynebacterium mastitidis*, we asked whether human-derived bacteria could do the same. Pairing sequencing and culturomics of healthy human conjunctiva from two geographically distinct cohorts, we define a stable core microbiome and identify *Corynebacterium bovis* as a consistent eye-resident commensal rather than a transient contaminant. In humans, *C. bovis* elicits a non-pathogenic response marked by IL-17 secretion by γδ T cells and circulating *C. bovis*-specific IgG, which suggests commensalism rather than pathology. Colonizing mice with human *C. bovis* strains from Pittsburgh or Miami reproduced this program and conferred protection against ocular infection with *Staphylococcus aureus*. Together, these findings identify *C. bovis* as a human-derived ocular commensal that can drive protective γδ T cell immunity, translating a mechanism previously defined only in mice into human relevance and a rational target against sight-threatening infection.

## INTRODUCTION

The human microbiome consists of communities of bacteria, viruses, fungi, and other microorganisms that can colonize sites throughout the body. In most cases, the microbiome supports immune development, provides colonization resistance against pathogens, aids host physiology and modulates inflammation. Dysbiosis of microbiomes can influence autoimmunity and infectious diseases. While most studies have focused on the intestinal microbiome and other high biomass sites, attention has moved towards low biomass sites like the ocular surface microbiome (OSM).

The OSM includes bacterial, fungal, and viral signatures; however, consistent identification of microbes has been difficult due to the paucibacterial nature of the ocular surface (1–3). Generally, OSM is isolated from the conjunctiva, or mucosal tissue, of the ocular surface with care taken to avoid contamination from the lid margins or skin adjacent to the ocular surface. Healthy conjunctival samples have approximately 150-fold less bacterial load than buccal mucosa as determined by 16S rDNA copies per host cell (1, 4). The reason for this disparity is largely attributed to limited nutrient availability, continuous tears flowing across the eye, high concentrations of tear anti-microbials like lysozyme, calprotectins, and others, and immunoglobulins like secretory IgA (sIgA).

The stability of the OSM remains a debated topic as recent deep sequencing studies fail to reach a consensus on core microbiome signatures. For example, a longitudinal study revealed that no conserved bacterial taxa were detected across 45 healthy individuals (4). That said, *coagulase-negative Staphylococcus*, *Propionibacterium*, and *Micrococcus* were present longitudinally in some individuals suggesting that a core ocular microbiome signature may be more influenced by individual nuances like factors related to personal decisions (sleep, extracurricular activities, smoking, etc), systemic disease, long-term medication, local environmental factors, dietary make-up, and many others. Widening the focus to the population level, more consistent signatures emerge. For example, *Corynebacteria*, *Acinetobacter*, *Pseudomonas*, *Staphylococci*, *Cutibacteria*, and *Streptococci* are consistently detected at the ocular surface. Although relative abundances vary across cohorts, repeated detection supports the notion that these bacteria should be included in the discussion of the core ocular microbiome. Furthermore, given the potential influence of many outside factors on OSM communities, it is not clear whether core OSM signatures are conserved across geographically distinct sites.

One challenge in examining these questions is that characterizing low-biomass communities is methodologically challenging due to limitations in both culture- and sequencing-based approaches. Culture dependent methods, which are ideal for isolating and purifying bacteria and fungi, may underestimate diversity as microbial recovery relies on appropriate growth media, environmental conditions, and incubation times. Conversely, sequencing based approaches can identify bacteria, viruses, and fungi; however, they may also detect DNA from non-viable organisms, may be contaminated by microbial signature in reagents, and influenced by sampling error. Given the limitations for individual methodologies, both approaches are required to comprehensively profile the ocular surface microbiome signatures.

Furthermore, to better understand the impact of the OSM on health, it is critical to define how it influences host physiology, including both immune and non-immune cells. Specifically, corneal and conjunctival epithelial cells can dynamically respond to microbes through the production and release of cytokines and chemokines. Additionally, the cornea and conjunctiva harbor immune cells that can mount responses affecting corneal integrity, wound healing, and resistance to infection. Ocular surface immune responses are particularly dynamic as previous studies have required topical application of antigen to stimulate the development of ocular mucosal immune responses (5–7). Similarly, mice topically treated with antibiotics or raised in germ free conditions have markedly reduced levels of secretory IgA (sIgA) and anti-microbials in the tears suggesting that local microbes promote immune activity at the ocular surface, and monitoring immune activity may help discern whether a microbe is a transient passerby or a true colonizer.

Here, we sought to address key gaps in knowledge regarding the composition and function of the OSM by comprehensively characterizing the healthy ocular surface microbiome and determining whether a core bacterial community exists at the ocular surface. We compared culture-independent sequencing techniques with culturomics from two geographically distinct clinical sites, Pittsburgh and Miami, to capture taxonomic breadth and viable bacteria at the ocular surface. Additionally, in order to study OSM function, we monitored whether one bacterial species found in samples from Pittsburgh and Miami, *Corynebacterium bovis*, stimulated physiological responses in human epithelial and immune cells. Complementing the human studies, we introduced human-derived ocular surface microbes into our mouse model of ocular surface colonization to evaluate colonization stability, host immune activation, and subsequent protection against infection with the Gram-positive bacterium *Staphylococcus aureus*. Our integrative and translational approach enabled a bridge between correlative data collection and mechanistic insight using cell culture and mouse models.

## Material and Methods

### Study Design and Recruitment of Healthy Subjects

This study was reviewed and approved by the University of Pittsburgh and University of Miami Ethics Committees (IRB: 23050083 and 20240425), and all procedures were conducted in accordance with the Declaration of Helsinki and the International Ethical Guidelines for Biomedical Research Involving Human Subjects. Prior to enrollment, each participant was fully briefed on the study’s objectives and procedures, and written informed consent was obtained from all individuals.

### Participant Recruitment and Eligibility

Participants were recruited from two clinical sites: the Department of Ophthalmology at UPMC Mercy Pavilion, Pittsburgh, Pennsylvania, and the Bascom Palmer Eye Institute, University of Miami, Florida. Eligibility required participants to be at least 18 years of age and willing to provide written informed consent. Exclusion criteria included age under 18 years, inability or unwillingness to provide consent, systemic or topical antibiotic use within the preceding three months, current use of prescription ocular medications, contact lens wear, systemic illnesses (e.g., Rheumatoid arthritis, Sjogren’s syndrome, Systemic lupus erythematosus, Diabetes mellitus, immunosuppression), symptoms of ocular surface disease (Dry Eye Questionnaire-5 (DEQ-5) ≥ 12), signs of ocular surface disruption (corneal staining > 2 out of 15) or Tear Break-Up Time (TBUT) <5 seconds, active ocular surface process (e.g., conjunctivitis, diabetic retinopathy, uveitis, glaucoma), anatomic abnormalities (e.g., pterygium, corneal edema, findings not consistent with age) or any ocular surgical procedure within the three months prior to enrollment.

### Sample collection

Ocular swab samples were collected from 50 individuals with a clinically healthy ocular surface (or Individuals not associated with active ocular diseases or abnormalities) each from two geographically distinct locations (Miami & Pittsburgh) for both nucleic acid extraction and culturomics-based analysis. Prior to swab collection, the ocular surface was anesthetized with one drop of Proparacaine Hydrocholoride ophthalmic solution 0.5% per eye. For both sequencing (right eye) and culturomics (left eye), conjunctival swabs were obtained by sweeping the lower conjunctival fornix three times using a counter-rotating technique relative to the direction of swab movement, taking care not to contact the lid margin. Swabs designated for nucleic acid sequencing (Zymo Research swabs; Fisher Scientific), collected from the right eye, were stored in DNA/RNA Shield Collection Tubes by snapping the swab tip directly into the prefilled collection device containing DNA/RNA Shield reagent (Zymo Research Corporation; catalog no. 50-125-1698, Fisher Scientific), in accordance with the manufacturer’s extraction protocol requirements. Negative template controls (NTCs) were generated by processing both unused swabs and swabs exposed to ambient air under identical conditions. All swabs were stored at −80°C prior to nucleic acid extraction.

### Sequencing

RNA extraction was performed using a commercially available kit and processed according to the manufacturer’s instructions (Zymobiomics DNA/RNA Miniprep Kit, Zymo Research) including an on-column RNase treatment. Negative controls included reagent blanks and positive controls included cells from a microbial community of known composition (ZymoBiomics Microbial Community Standards; Zymo Research). All samples were first spiked with 20ul of ZymoBIOMICS Spike-in Control II prior to extraction. Approximately 1 ng of extracted total RNA from each sample was reverse transcribed in an RT reaction using Superscript IV VILO (ThermoFisher) as follows. gDNA was first digested in a 10ul volume using the ezDNase enzyme at 37°C for 2 minutes, followed by the addition of SuperScript IV VILO master mix and nuclease free water to a volume of 20ul. The reaction was incubated at 25°C for 10 minutes, 50°C for 10 minutes, 85°C for 5 minutes. PCR was performed on 5ul cDNA from each sample, in a 25 μL volume using Q5 HS High Fidelity polymerase (New England BioLabs, Ipswich, MA). Cycles were 98°C for 30 seconds, then 30 cycles of 98°C for 10 seconds, 57°C for 30 seconds, and 72°C for 30 seconds, followed by a single 72°C incubation for 2 minutes. PCR products were purified with AMPure XP beads (Beckman Coulter, Indianapolis, IN), with a ratio of 0.8:1 (beads:DNA). Purified DNA was then quantified using a Qubit fluorimeter (Life Technologies, Grand Island, NY). Final pooling used 40 ng of each purified band or 20ul for any sample with low concentration. Two subsequent purifications were performed using MinElute PCR purification kit (Qiagen AMPure XP beads (0.8:1)) and Purelink PCR Purification Kit (Life Technologies). Quantification of the final pool was performed on the Qubit fluorimeter before sequencing. Preparation of the sequencing pool was performed according to Illumina’s protocol (Illumina, Inc., San Diego, CA); immediately after the initial dilution to 20pM, an additional incubation at 95°C for 2 minutes was performed. A final dilution was performed to 6pM + 25% PhiX control (Illumina), and Illumina MiSeq 500 cycle V2 kits were used for sequencing.

### Bioinformatics

Sequences were deconvolved on the MiSeq. Processing was performed using an in house sequence quality control pipeline. Dust low complexity filtering, quality value trimming, trimming of primers used for 16S rRNA gene amplification, and minimum read length filtering was performed. Contigs were created by merging forward and reverse reads. Processing occurred through our institution’s mothur based 16S clustering and sequence annotation pipeline (mothur V1.48.2, silva V138.2). Sequence taxonomic classifications were developed using the Ribosomal Database Project (RDP) Naive Bayesian Classifier with the Silva reference database.

Spiked in Zymo controls (*Truepera, Imtechella, and Allocbacillus*) and potential contaminants (*Escherichia Shigella, Salmonella, and Enterobacteriaceae_uncl*) also found in the RT reagents were removed. Computational distribution-based background subtraction was performed on the taxonomic profiles between the kit extraction and RT assay negative controls and the profiles generated from the experimental samples (8).

### Associations between Ocular Microbiome and Selected Covariates

Three complementary methods were used to examine whether characteristics of the ocular surface microbiome were associated with smoking history, sex, age, study location, systemic allergies, antihistamine use, and artificial tear use. Smoking was assessed using two variables to distinguish current smoking from prior exposure: participants were classified as current smokers, former smokers, or never smokers. This allowed us to examine potential effects of active smoking separately from those associated with a history of smoking. Study location distinguished participants recruited in Pittsburgh, Pennsylvania, from those recruited in Miami, Florida.

#### Distance-Based

The distance-based methodology utilizes PERMANOVA through the R package vegan and the adonis2() function. PERMANOVA associates the inter-sample distance (via the Manhattan distance) with covariates. LocationID was significantly associated with a Cohen’s effect size of 0.02 (small effect) and a p-value of 0.012. Hierarchical clustering with multinomial logistic regression (HCAMLR) (9) a distance-based method that first performs hierarchical clustering across the samples, then iteratively cuts the tree into k clusters and then uses multinomial logistic regression to associate subject covariates with the resultant k clusters was applied.

#### Distribution-Based

Distribution-based analyses utilized the Tail and Shannon diversity index to quantify the compositional diversity within each sample. Although the Shannon diversity index is more commonly accepted to quantify microbial diversity, capturing the most abundant taxa present and how evenly they are distributed. However, it can miss compositional differences that are really being driven by rarer, low-abundance organisms. However, The Tail statistic is more sensitive to differentiating microbiome samples, since it places a greater emphasis on the abundance distribution’s less dominant components (10). This makes it more sensitive than the Shannon index for detecting differences between samples that are driven by subtle shifts in rare taxa rather than dominant ones. Using both metrics together allowed us to capture diversity differences across the full abundance spectrum from dominant to rare taxa rather than relying on a single measure that may overlook biologically meaningful variation.

### Culturomics - High-Throughput Culturing

Conjunctival swabs were collected from the left eye of each healthy subject using BD BBL™ CultureSwab EZ™ (Fisher Scientific) and immediately submerged in sterile tubes containing 1ml of thioglycolate broth. To ensure standardized processing times across geographically distinct collection sites, samples obtained at the University of Pittsburgh were stored on ice for 24 hours prior to processing, while samples collected in Miami were placed on ice and shipped overnight before processing.

To facilitate release of bacteria from the swab into the broth, tubes were vigorously vortexed for 5 minutes. Subsequently, 100 µL of the primary thioglycolate broth was inoculated into two parallel sets of six enrichment broths: (1) Brain Heart Infusion supplemented with 5% sheep blood (BHI+5%), (2) Cooked Meat Medium (CMM), (3) de Man, Rogosa and Sharpe broth (MRS), (4) Reinforced Clostridial Medium (RCL), (5) Columbia broth supplemented with 5% sheep blood (CLA+5%), and (6) Luria-Bertani broth supplemented with Tween-80 and 5% sheep blood (LBT+5%). Given the inherently aerobic nature of the ocular surface environment, all enrichment cultures were incubated under two conditions, stationary aerobic and shaking at 150 rpm at 37°C for 48 hours.

Following enrichment, cultures were subcultured onto corresponding solid media: BHI+5% broth → BHI+5% agar; CMM broth → CMM agar; MRS broth → MRS agar; RCL broth → RCL agar; LBT+5% broth → LBT+5% agar; and CLA+5% broth → Chocolate agar (CHA). All plates were incubated under aerobic conditions at 37°C for 24–48 hours. To monitor environmental contamination, swabs were waved openly within the collection room and processed in parallel as negative controls. Broth-only controls were likewise included throughout the expansion process to verify the absence of exogenous microbial contamination introduced during handling.

### 16S rRNA Gene Sequencing and Taxonomic Classification

Following 24–72 hours of incubation, bacterial colonies were identified by 16S rRNA gene sequencing, as previously described (11). Sequences sharing less than 98% similarity with entries in reference databases were designated as putative novel bacterial species, while those sharing less than 95% similarity were classified as putative novel bacterial genera.

### Immune profiling using flow cytometry

Human conjunctival cells were collected using the Eyeprim device (12). Samples were collected from the temporal bulbar conjunctiva following topical anesthesia with 0.5% proparacaine ophthalmic solution. After 2 minutes, participants were instructed to look nasally, and the EyePrim membrane was applied perpendicular to the temporal bulbar conjunctiva while avoiding contact with the eyelids, eyelashes, and surrounding skin. The membrane was held in contact with the ocular surface for 10 seconds and then transferred into sterile 1.5 mL tubes containing Roswell Park Memorial Institute (RPMI) media + 10% fetal bovine serum (FBS). Samples were maintained at 4°C until processing. For flow cytometric analysis, samples were transferred to Fluorescence-Activated Cell Sorting (FACS) tubes containing FACS buffer (0.1% Sodium Azide, 1% FBS, 1X Phosphate-Buffered Saline (PBS)) centrifuged at 1400 RPM for 4 minutes at 4°C, and stained with an antibody cocktail containing CD4 AF488, CD3 PECy7, CD45 BV510, γδ TCR PE, CD8 BV42, αβ TCR APC, CD19 BV605, CD16 APC Fire750, CD15 AF700, CD49d BV71, CD56 PE-Dazzle 594, and CD11b BV650. Following incubation at 4°C protected from light, samples were washed, fixed with 4% paraformaldehyde (PFA), and analyzed by flow cytometry.

#### Heat Inactivation of *Corynebacterium* Strains

Human-derived *Corynebacterium bovis* strains isolated from Miami (*mC. bovis*); Pittsburgh (*pC. bovis*), and *C. mastitidis* (Mice strain; AS1) were cultured in Luria-Bertani broth supplemented with Tween-80 (LBT) and with 100 μg/mL fosfomycin for Corynebacterium selection were grown for 12–18 hours at 37°C under orbital shaking condition. Bacterial density was quantified spectrophotometrically at an optical density of 600 nm (OD□□□). Cultures were then centrifuged at 8,000 × g for 5 minutes, and the pellet was washed with 1X PBS. Bacteria were resuspended in 1x PBS at the concentration required for downstream assays and heat-inactivated at 65°C for 30 minutes.

#### Characterizing genomic differences between *mC. bovis* and *pC. bovis*

To characterize the genomic and functional differences between the two human-derived Corynebacterium strains (*mC. bovis* and *pC. bovis*), genomic DNA was extracted using a DNA extraction kit (Takara) according to the manufacturer’s instructions. DNA quality and concentration were assessed using a NanoDrop spectrophotometer. Purified genomic DNA was then submitted to Plasmidsaurus (USA) for whole-genome sequencing on the Oxford Nanopore platform. The contigs were assembled and annotated using Comprehensive Genome analysis and then compared with Comparative Systems and Proteome comparisons from PATRIC(13).

#### *S. aureus* co-cultures with *C. bovis* with and without Xanthine

To test if Xanthine Phosphoribosyltransferase activity could affect the pathogenicity of *S. aureus* we grew both strains of *C. bovis* in triplicate to log phase in Davis minimal media containing 0.5% corn oil and Casamino acids with and without 2mM of Xanthine. We chose this concentration as previous studies have shown elevated xanthine levels in the tears of keratitis patients reached 2mM (14). We then diluted each culture to 0.5 OD_600_ with more media and filter sterilized each culture. The conditioned media from each culture was then divided into four 500μL wells on a 48 well plate; 2 wells were inoculated with 10μL of an overnight culture of *S. aureus* and 2 wells were left as blank controls to ensure the *Corynebacteria* were properly sterilized. Unused media with and without *S. aureus* were used as controls. The growth was measured over 36 hours with the Spectramax 3 plate reading spectrophotometer and the growth curves and calculations of area under the curve were completed using GraphPad Prism software.

### Isolation and Stimulation of Peripheral Blood Mononuclear Cells (PBMCs)

To examine the immunostimulatory potential of ocular surface bacteria on human peripheral blood mononuclear cells (PBMCs), peripheral blood was obtained from 12 healthy ocular surface volunteers; six from Pittsburgh and six from Miami. PBMCs were isolated using the ACCUSPIN System with Histopaque-1077 density gradient separation medium (Millipore Sigma). Following gradient centrifugation, the PBMC containing layer and plasma fraction were collected and stored at −80°C for downstream analysis.

Isolated PBMCs were resuspended in stimulation medium consisting of DMEM supplemented with 10% fetal bovine serum (FBS), sodium pyruvate, HEPES, non-essential amino acids (NEAA), and glutamine (all components diluted 1:100). Co-culture assays were performed in 96-well plates at a density of 1x10□ bacteria and 5x10□ PBMCs/ml. Control wells contained PBMCs in medium alone; experimental wells contained PBMCs co-cultured with bacteria in stimulation medium (*mC. bovis, pC. bovis*, or *C. mastitidis* (Mice strain; AS1). PBMCs stimulated with anti-CD3 antibody (aCD3; 6 µg/ml) served as positive control. All co-cultures were incubated at 37°C in 5% CO□ for 72 hours.

For phenotypic characterization of PBMC activation, cells were stained with the following fluorochrome-conjugated antibodies (BioLegend): CD3-PECy7, CD45-BV510, CD8α-BV421, CD4-AF488, γδ TCR-PE, TCRβ-APC, and CD69-PerCP/Cyanine5.5, and incubated for 30 minutes. Cells were subsequently washed, fixed, and stained intracellularly with Ki67-AF700 (BioLegend) for 30 minutes. Following a final wash, cells were resuspended in the FACS buffer and acquired by flow cytometry.

### Bacterial-Specific Cytokine Analysis

To evaluate cytokine secretion with particular focus on IL-17 production; PBMC co-cultures were incubated in antibiotic-free conditions and sampled at 2, 24, 48, and 72 hours post-stimulation at 37°C in 5% CO□. Cell culture supernatants were collected at each time point and stored at −20°C until analysis. Cytokine concentrations were quantified by enzyme-linked immunosorbent assay (ELISA) according to the manufacturer’s instructions (R&D Systems).

### Quantification of Inflammatory Cytokines in Human Corneal Epithelial Cell

To assess the epithelial cytokine response to commensal *Corynebacterium* stimulation, human corneal epithelial (HCLE) cells were co-incubated with heat-killed preparations of *mC. bovis*, *pC. bovis*, and *C. mastitidis* for 24, 48 and 72 hours. Following incubation, cell culture supernatants were carefully collected and stored at −80°C until analysis. Secreted concentrations of IL-6, IL-8, and CXCL10 were quantified using a multiplexed bead-based immunoassay (LEGENDplex Human Inflammation Panel, BioLegend), performed in accordance with the manufacturer’s instructions. All experimental conditions were performed in triplicate.

### Bacteria-Specific Antibody Responses

To assess humoral immune responses directed against ocular surface bacteria, plasma obtained from healthy subjects was serially diluted to final concentrations of 1:200, 1:500, 1:1000, and 1:2000 in FACS buffer. Diluted plasma samples were incubated with *mC. bovis, pC. bovis*, or *C. mastitidis* (Mice strain; AS1) at a concentration of 1x10□ CFU/ml for 30 minutes. Following incubation, samples were stained with anti-human IgG antibodies conjugated to PE and APC, respectively (both at 1:500 dilution), for 30 minutes. Samples were washed with FACS buffer, resuspended, and acquired on a CytoFlex flow cytometer (Beckman Coulter). Data was analyzed using CytExpert software.

### In Vivo Studies

#### Specific Pathogen-Free and Germ-Free Mouse Models

All animal procedures were approved by the University of Pittsburgh Institutional Animal Care and Use Committee (IACUC protocol number: 23063056) and conducted in compliance with the ARVO Statement for the Use of Animals in Ophthalmic and Vision Research. Germ-free (GF) C57BL/6J mice were bred and maintained in flexible film gnotobiotic isolators at the Gnotobiotic Animal Core Laboratory, University of Pittsburgh. Age-matched specific pathogen-free (SPF) C57BL/6J mice harboring a conventional complex microbiota were purchased from The Jackson Laboratory (Bar Harbor, ME) and housed under SPF conditions at UPMC Mercy Pavilion. GF mice received autoclaved lab diet, and all animals had unrestricted access to water, which was sterilized for GF cohorts. For this study, 12 male and 12 female GF mice and 15 female SPF mice at 6 weeks of age were used to assess colonization by human-derived ocular surface bacteria.

#### Colonizing Both SPF and GF mice with Human Specific *C. bovis*

Both SPF and GF mice were inoculated ocularly with human-derived *Corynebacterium bovis* strains isolated from Miami (*mC. bovis*) and Pittsburgh *(pC. bovis*). Inoculations were administered every other day for a total of three doses at a concentration of 5x10□ CFU per eye.

#### Tissue Harvest and Conjunctival Cell Isolation

Three weeks post-inoculation, both SPF and GF mice were sacrificed by cardiac perfusion, and conjunctival tissue was isolated as previously described (15, 16). Briefly, conjunctival tissue from one eye was excised and transferred to 500 µL of 1x PBS for bacterial CFU enumeration. Tissue from the contralateral eye was finely minced and enzymatically digested in collagenase solution at 37°C for one hour. Following digestion, the tissue suspension was passed through a Falcon cell strainer (Corning, USA) to obtain a single cell suspension.

#### Immune Profiling of Conjunctival Cells and Assessment of *C. bovis* Induced Immune Activation In Vivo

Isolated conjunctival cells were either directly stained for comprehensive immunophenotyping or stimulated with lymphocyte activation cocktail (1:500 dilution) for 4 hours at 37°C prior to staining. Following stimulation, cells were stained with the following fluorochrome-conjugated antibodies: TCRαβ-BV421, CD90.2-BV605, γδTCR-AF488, Vγ2 TCR-PerCP eF700, CD4-BV785, CD8-APC AF780, CCR6-PerCP Fire 780, Vγ4 TCR, and IL-17A. All samples were acquired and analyzed using a spectral ID7000 Flow Cytometer (Sony Biotechnology).

#### CFU Enumeration in Mice Colonized with *C. bovis*

Conjunctival tissue collected in 1x PBS was homogenized and plated directly onto Luria-Bertani agar supplemented with Tween-80 (LBT) and fosfomycin (100 µg/ml) for selective enumeration of Corynebacterium species. Plates were incubated at 37°C for 24 - 48 hours, after which colony-forming units (CFUs) were counted per eye to quantify the extent of ocular colonization. Representative colonies were subsequently subcultured and subjected to 16S rRNA gene sequencing to confirm the identity of the recovered isolates as human-derived *Corynebacterium bovis* strains.

#### Pathogenic *S. aureus* challenge in mice colonized with Corynebacterium

A clinical *S. aureus* (Methicillin-resistant Staphylococcus aureus (MRSA))strain obtained from the Charles T. Campbell Eye Microbiology Laboratory was cultured overnight in LB broth at 37°C with shaking. Cultures were centrifuged at 8,000 × g for 5 minutes at 25°C, and the resulting bacterial pellets were washed and resuspended in sterile phosphate-buffered saline (PBS). Bacterial concentrations were confirmed prior to inoculation, and working suspensions were adjusted to 1×10□ CFU/eye in a 5 µl inoculum volume.

#### Colonization and Corneal Infection Model

Mice were pre-colonized with human-derived *C. bovis* strains (*mC. bovis* and *pC. bovis*; 5×10□ CFU per eye) three weeks prior to pathogen challenge, allowing sufficient time for stable ocular surface colonization. At the time of challenge, mice were anesthetized via intraperitoneal injection of a ketamine (100 mg/kg) and xylazine (10 mg/kg) mixture. To disrupt the corneal epithelial barrier and establish a standardized infection model, 8–15 vertical and horizontal scratches were made across the central corneal epithelium using a dulled 30-gauge needle. Immediately following scarification, the right eye of each mouse was inoculated with 5 µl of *S. aureus* suspension (1×10 CFU/eye). Scratch-only eyes receiving 5 µl of sterile 1×PBS served as negative controls.

Ocular disease severity was assessed at 24 and 48 hours post-infection by a scientist masked and scored using a Zeiss Stemi 305 stereomicroscope. Corneal pathology was graded on a 0–4 scale adapted from Girgis et al., **0** — Clear cornea; no opacity; **1** —Readily detectable but mild corneal opacity; **2** — Opacity partially covering the corneal surface over the pupil; **3** — Dense opacity covering the entire corneal surface over the pupil; **4** — Moderate to dense opacity covering the entire corneal surface with corneal erosion.

#### Pathogenic *S. aureus* quantification

Following the clinical scoring, mice were euthanized and whole eyes were enucleated aseptically. Each eye was placed in a 2 ml tube containing 500 µL of sterile PBS and 1.4 mm ceramic beads (Fisher Scientific) and homogenized using an MP FastPrep-24 tissue homogenizer. A 100 µl aliquot of each homogenate was serially diluted 1:1000 in PBS and plated on LB agar (Fisher Scientific). Plates were incubated at 37°C for 24 hours, after which bacterial colonies were enumerated to determine ocular CFU burden.

### Statistical Analysis

All statistical analyses were performed using GraphPad Prism software (version 11.01). For normally distributed data, comparisons among three or more groups were conducted using one-way ANOVA followed by Dunnett’s multiple comparisons test to assess individual group differences relative to a reference condition. For non-normally distributed data, the Kruskal-Walli’s test was applied as the non-parametric equivalent for comparisons across three or more independent groups. Statistical significance was defined as p-value of less than 0.05.

## RESULTS

The mean age of participants was 33.46 ± 12.52 years (Miami 28.14 ± 7.80; Pittsburgh 38.78 ± 14.07); 67.0% self-identified as female, 76.0% as White, and 14.0% as Hispanic (Table 1). None reported clinically significant ocular surface symptoms on standardized questionnaires (Ocular Surface Disease Index 2.62 ± 3.95; 5-Item Dry Eye Questionnaire 2.15 ± 2.26), and tear parameters were healthy (Table 1).

**Table 1:**
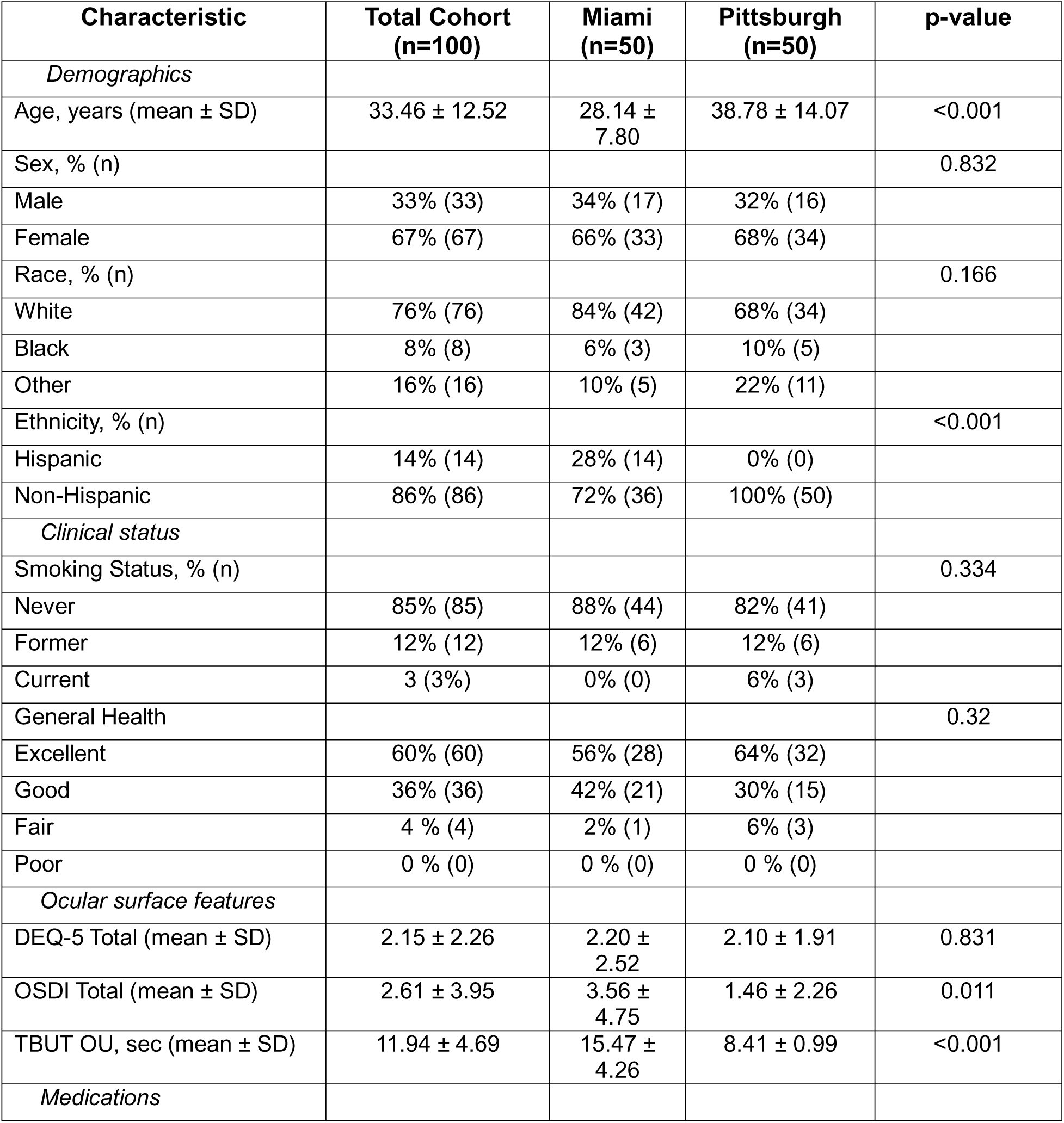
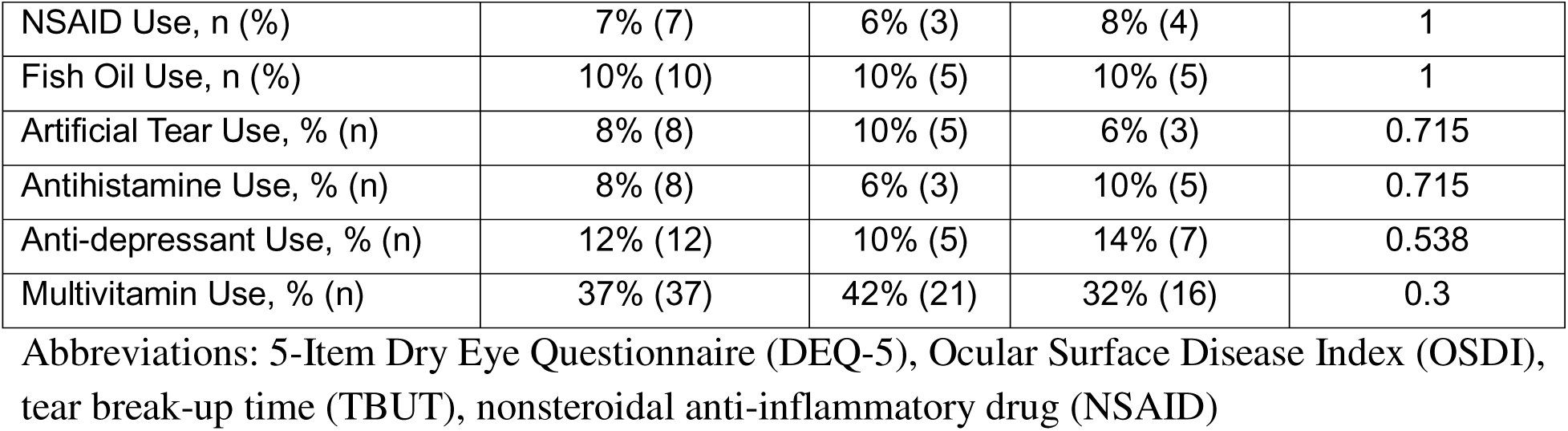
Study Population Demographics.

| **Characteristic** | **Total Cohort (n=100)** | **Miami (n=50)** | **Pittsburgh (n=50)** | **p-value** |
|---|---|---|---|---|
| *Demographics* |  |  |  |  |
| Age, years (mean ± SD) | 33.46 ± 12.52 | 28.14 ± 7.80 | 38.78 ± 14.07 | <0.001 |
| Sex, % (n) |  |  |  | 0.832 |
| Male | 33% (33) | 34% (17) | 32% (16) |  |
| Female | 67% (67) | 66% (33) | 68% (34) |  |
| Race, % (n) |  |  |  | 0.166 |
| White | 76% (76) | 84% (42) | 68% (34) |  |
| Black | 8% (8) | 6% (3) | 10% (5) |  |
| Other | 16% (16) | 10% (5) | 22% (11) |  |
| Ethnicity, % (n) |  |  |  | <0.001 |
| Hispanic | 14% (14) | 28% (14) | 0% (0) |  |
| Non-Hispanic | 86% (86) | 72% (36) | 100% (50) |  |
| *Clinical status* |  |  |  |  |
| Smoking Status, % (n) |  |  |  | 0.334 |
| Never | 85% (85) | 88% (44) | 82% (41) |  |
| Former | 12% (12) | 12% (6) | 12% (6) |  |
| Current | 3 (3%) | 0% (0) | 6% (3) |  |
| General Health |  |  |  | 0.32 |
| Excellent | 60% (60) | 56% (28) | 64% (32) |  |
| Good | 36% (36) | 42% (21) | 30% (15) |  |
| Fair | 4 % (4) | 2% (1) | 6% (3) |  |
| Poor | 0 % (0) | 0 % (0) | 0 % (0) |  |
| *Ocular surface features* |  |  |  |  |
| DEQ-5 Total (mean ± SD) | 2.15 ± 2.26 | 2.20 ± 2.52 | 2.10 ± 1.91 | 0.831 |
| OSDI Total (mean ± SD) | 2.61 ± 3.95 | 3.56 ± 4.75 | 1.46 ± 2.26 | 0.011 |
| TBUT OU, sec (mean ± SD) | 11.94 ± 4.69 | 15.47 ± 4.26 | 8.41 ± 0.99 | <0.001 |
| *Medications* |  |  |  |  |
| NSAID Use, n (%) | 7% (7) | 6% (3) | 8% (4) | 1 |
| Fish Oil Use, n (%) | 10% (10) | 10% (5) | 10% (5) | 1 |
| Artificial Tear Use, % (n) | 8% (8) | 10% (5) | 6% (3) | 0.715 |
| Antihistamine Use, % (n) | 8% (8) | 6% (3) | 10% (5) | 0.715 |
| Anti-depressant Use, % (n) | 12% (12) | 10% (5) | 14% (7) | 0.538 |
| Multivitamin Use, % (n) | 37% (37) | 42% (21) | 32% (16) | 0.3 |

Abbreviations: 5-Item Dry Eye Questionnaire (DEQ-5), Ocular Surface Disease Index (OSDI), tear break-up time (TBUT), nonsteroidal anti-inflammatory drug (NSAID)

### Sequencing Conjunctival Samples Identifies *Staphylococcus*, *Corynebacterium*, and *Lawsonella* as core members of the OSM

The commonly shared taxa of the ocular microbiome, i.e. the “core”, was defined using a 2-parameter model previously introduced to examine the cohort whose samples were sequenced for the Human Microbiome Project (HMP) (10). A taxon enters the core only if it meets two criteria: a minimum relative abundance, and a minimum proportion of subjects exceeding that abundance (its ubiquity). This cleanly separated the core from the rest of the microbiome, and we plotted the non-core taxa alongside it to show how well the two groups separate (Figure 1A). From the active taxa classified from 16S rRNA (as opposed to 16S rRNA genes from DNA), those taxa identified at an abundance exceeding 0.01 (i.e., log10 (.01) =-2), and ubiquity exceeding 60% of the cohort, the core ocular microbiome consists of *Corynebacterium*, *Lawsonella* and *Staphylococcus*. Notably, minor adjustments to the cutoff for abundance yields the possibility that *Actinomycetota* can be included in the core OSM.

**Figure 1.**
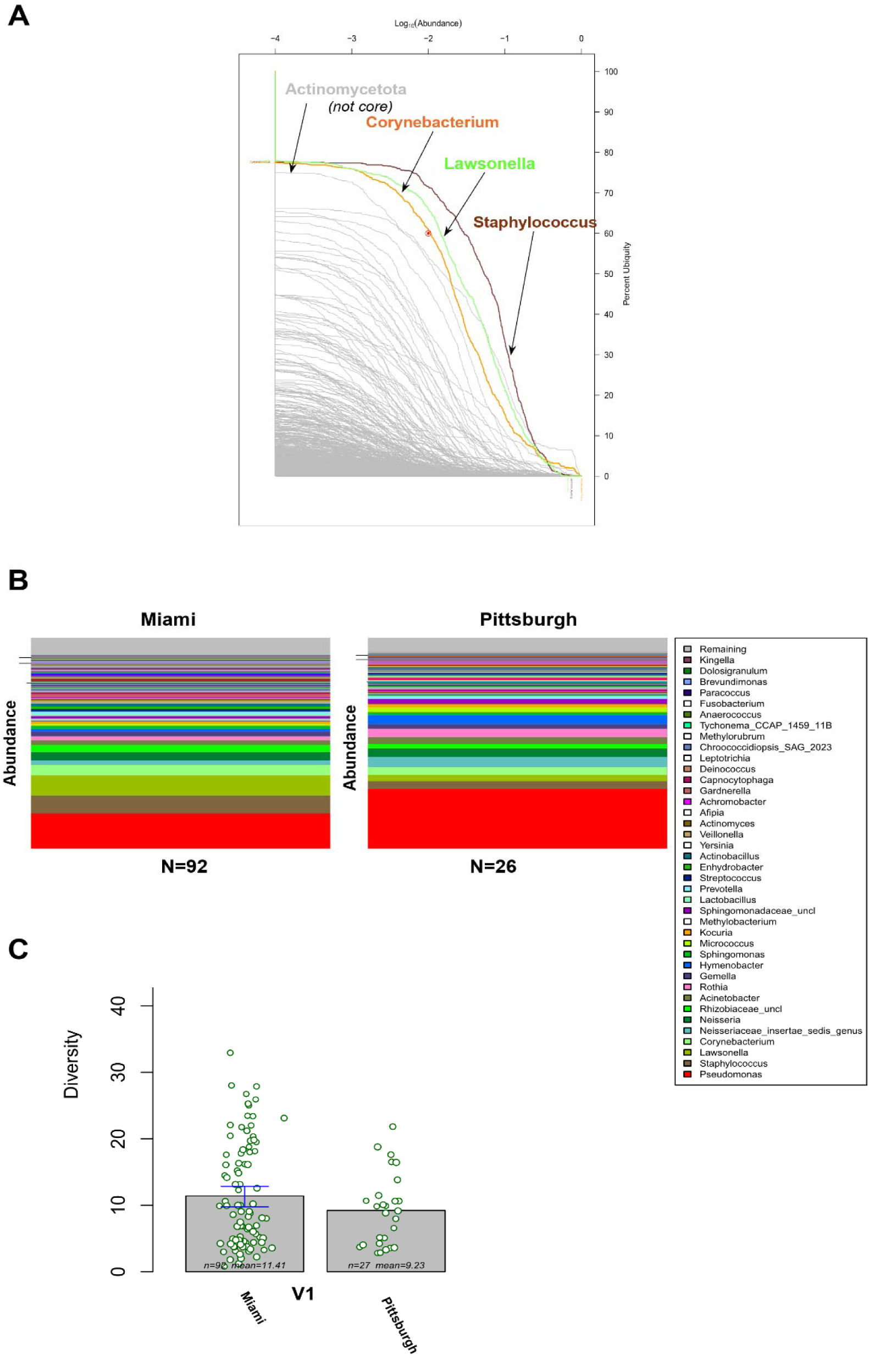
Sequencing of conjunctival samples identifies *Staphylococcus*, *Corynebacterium*, and *Lawsonella* as core members of the ocular surface microbiome (OSM). **(A)** Ubiquity-abundance plot showing the percent ubiquity as a function of log₁₀ relative abundance for all bacterial genera identified across conjunctival samples. Each gray line represents an individual genus; colored lines highlight *Corynebacterium* (orange), *Lawsonella* (green), and *Staphylococcus* (dark red) as high-ubiquity, high-abundance taxa comprising the core OSM. **(B)** Stacked bar plots depicting the relative abundance of bacterial genera identified in conjunctival samples collected from the Miami (N = 92) and Pittsburgh (“Pitt,” N = 26) sites. Each color represents a distinct bacterial genus, as indicated in the legend; “Remaining” denotes the cumulative abundance of low-abundance taxa not individually color-coded. **(C)** Bacterial diversity (Shannon diversity index) of conjunctival samples from healthy ocular surface participants at the Miami (HCF) and Pittsburgh (HCP) sites. Each dot represents an individual sample; bars represent group means with (SEM/95% CI).

Given the notable influences that environmental factors have on the make-up of microbiome at the ocular surface and at other sites in the body (17–19), we sought to investigate whether there were significant differences between the OSM of samples collected in Pittsburgh and Miami. When analyzing samples by location, we find that bacterial genera are generally similar (Figure 1B) and the level of diversity in the represented taxa in OSM samples is not significantly different between the Pitt and Miami locations (Figure 1C). *Pseudomonas* made up a sizeable part of the microbiome signatures in some Miami and Pitt samples; however, it was not considered part of the core OSM signature (Figure 1A & B) due to limited ubiquity across many samples (Supplemental Figure 1A). Given its propensity to contaminate low biomass samples (20, 21) and its detection in only a small subset of healthy subjects (Supplemental Figure 1A), we asked why some subjects harbored *Pseudomonas* and others did not. No covariate in Table 1 tracked with the Pseudomonas cluster, suggesting these samples were not outliers from sampling bias. We therefore judged that Pseudomonas-containing samples may not reflect the true core, but we retained this signature to ensure rigorous analysis of the data.

Despite similar diversity between locations, location itself was a significant determinant of the OSM signature (Supplemental Figure 1A). The healthy eye appears to support a consistent number of distinct microbes, but geography influences which bacteria fill those slots. Using the metadata in Table 1, we then asked whether other covariates shape the OSM, and found that location, sex, artificial tear use, and systemic allergies each significantly influenced the global make-up of the 35 most abundant taxa (Supplemental Figure 1B).

Additionally, we found specific taxonomic associations with covariates that did not necessarily alter the general ocular microbiome signature. Upon further investigation of these specific taxonomic associations, we found that Acinetobacter was lost in aged subjects (Supplemental Figure 1C) and was elevated in subjects with systemic allergies (Supplemental Figure 1D). Females had significant reductions in *Staphylococcus* (Supplemental Figure 1E). Notably, smoking, which can also dramatically influence microbiomes, was connected to elevated levels of *Rosemonas* (Supplemental Figure 1F) compared to non-smokers. Finally, Pittsburgh had marked losses of *Gardnerella*, *Actinobacillus*, *Veillonella*, and *Lactobacillus* compared to Miami. Conversely, *Kocuria* was elevated in Pittsburgh samples compared to Miami samples (Supplemental Figure 1G). Other taxa trended with these same covariates without reaching significance (Supplemental Figure 1C-I). In sum, the abundance of specific taxa tracked with several participant and exposure-related characteristics.

### *Staphylococcus* and *Corynebacterium spp.* can reliably be isolated from human conjunctiva

Given the possibility of genetic material coming from dead or dying microbes skewing sequencing results, we sought to isolate and identify live bacteria from the ocular surface to further support the notion that bacteria may be living at the ocular surface. Using a culturomics-based approach that was previously described (11, 22), we detected microbial growth in 47% of the eye swabs collected, recovering a total of 59 bacterial isolates from eye swabs of healthy individuals in Pittsburgh and Miami. A total of 42 isolates (71%) came from Pittsburgh, with the remaining 17 (29%) from Miami.

Within this analysis, *Staphylococcus epidermidis* and *Corynebacterium bovis* emerged as the most prevalent species overall, found in 47% (28/59) and 17% (10/59) of isolates, respectively (Figure 2A). At the genus level, *Staphylococcus* and *Corynebacterium* dominated the ocular microbiome signature at both sites (Figure 2B). *Staphylococcus* was present in 64.7% (11/17) of Miami isolates and 43% (18/42) of Pittsburgh isolates, while *Corynebacterium* accounted for 35% (6/17) and 19% (8/42) of isolates from the two sites, respectively. At the species level, *S. epidermidis* was the leading staphylococcal species at both locations (Miami: 59%; Pittsburgh: 43%), and *C. bovis* was the most common corynebacterial representative at each site (Miami: 23.5%; Pittsburgh: 14%). Beyond these dominant species, the two sites differed considerably. Miami isolates were restricted to *C. mastididis* and *S. hominis*, whereas Pittsburgh subjects harbored a richer diversity that included *S. aureus*, *S. warneri*, *S. capitis*, *S. saprophyticus*, *C. mastitidis*, *C. macginleyi*, and *C. lowii* (Figure 2B–C). Overall, dominant taxa were distributed consistently between sites, indicating a broadly similar live microbial profile that corroborates our sequencing results.

**Figure 2.**
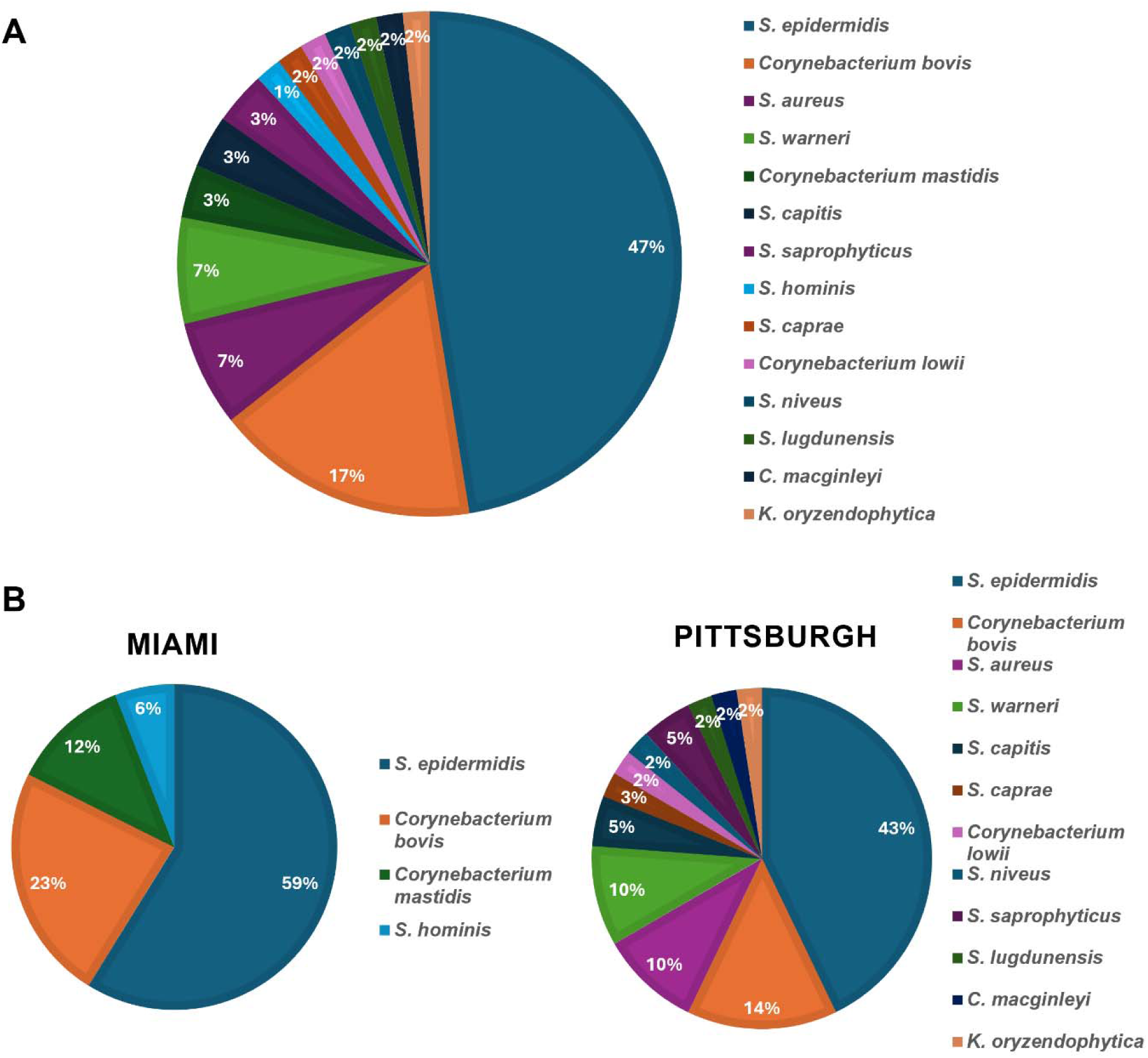
*Staphylococcus* and *Corynebacterium* spp. can reliably be isolated from human conjunctiva. (A) Pie chart depicting the relative proportion of bacterial species cultured from human conjunctival swabs across the study sites (Maimi and Pittsburgh) combined. *Staphylococcus epidermidis* (47%) and *Corynebacterium bovis* (17%) were the most frequently isolated species, followed by other *Staphylococcus* and Corynebacterium spp. (<7%). (B) Pie charts depicting the relative proportion of bacterial species cultured from conjunctival swabs collected at the Miami and Pittsburgh sites, shown separately. At the Miami site, isolates were limited to four species: *S. epidermidis* (59%), *C. bovis* (23%), *C. mastidis* (12%), and *S. hominis* (6%). At the Pittsburgh site, a broader diversity of species was recovered, including *S. epidermidis* (43%), *C. bovis* (14%), *S. aureus* (10%), *S. warneri* (10%), *S. capitis* (5%), *C. mastidis* (5%).

### Comparing Genome Sequences of two isolates of *C. bovis*

From our culturomics approach, we isolated two strains of *C. bovis*, one from the Pittsburgh site (*pC. bovis*) and one from the Miami site (*mC. bovis*) (Figure 2A). Whole-genome sequencing on ONT and assembly of the two strains yielded genomes of 2.7 Mbp (2 contigs, N50 = 7,750 bp; GC content = 72.71%) and 2.6 Mbp (2 contigs, N50 = 3,938 bp; GC content = 72.89%), respectively (Figure 3A). Circular genome maps generated in Bakta showed coding sequencing, tRNA, rRNA, and non-coding RNA genes on both strands alongside GC content and GC skew. Genome size differed only modestly between the strains (0.1 Mbp, or 4%) with no major change in gene organization.

**Figure 3.**
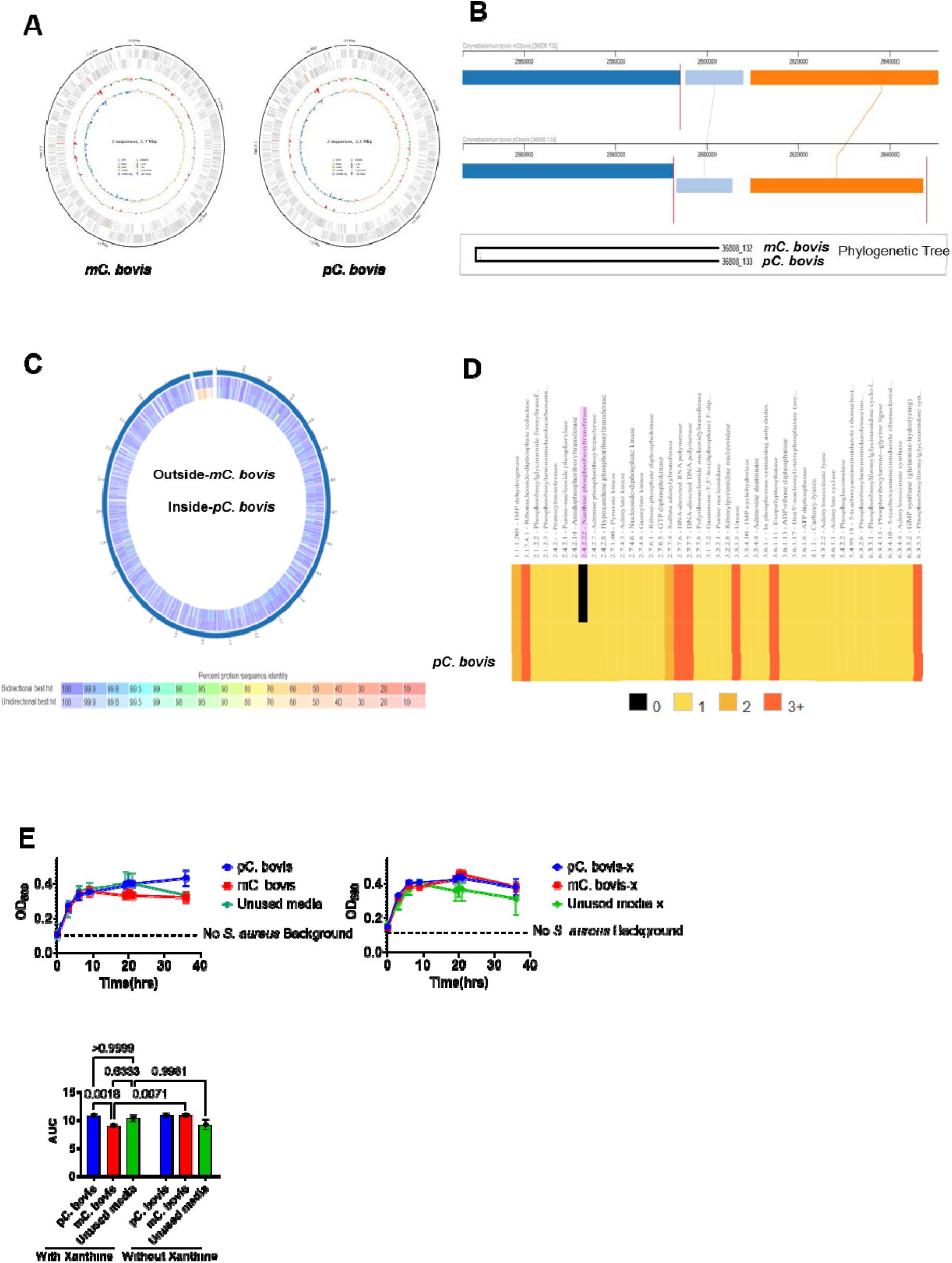
Genomic and functional differences between the human-derived *C. bovis* strains. **(A)** Circular genome maps of *mC. bovis* and *pC. bovis* generated using Bakta, depicting coding sequences (CDS) on the forward and reverse strands, tRNA, rRNA, and non-coding RNA genes, GC content, and GC skew. Both genomes consist of two sequences, with total assembly sizes of 2.7 Mbp (*mC. bovis*) and 2.6 Mbp (*pC. bovis*). **(B)** Whole-genome alignment of *mC. bovis* and *pC. bovis* generated using the BV-BRC Genome Comparison Viewer, illustrating a high degree of synteny across the shared genomic backbone. A structural rearrangement is highlighted within the ∼2,550,000– 2,650,000 bp region, in which a genomic segment (light blue block) is repositioned relative to an adjacent block (orange) between the two strains; red vertical lines indicate contig boundaries. Core-genome phylogenetic tree (bottom) showing the evolutionary relationship between *mC. bovis* and *pC. bovis*. **(C)** Circular proteome comparison plot generated in BV-BRC depicting bidirectional BLAST best-hit percent protein sequence identity between *mC. bovis* (outer track) and *pC. bovis* (inner track). A subset of loci displayed reduced identity (70–90%), indicating localized strain-specific divergence or loss of a small number of genes. **(D)** Heatmap depicting predicted enzyme copy number across genes associated with the purine metabolism pathway in *mC. bovis* and *pC. bovis*, generated using BV-BRC comparative pathway analysis. Copy number is indicated by color: 0 (black), 1 (yellow), 2 (orange), and 3+ (red). Xanthine phosphoribosyltransferase (highlighted in pink) is the enzyme in this pathway showing a presence/absence difference, being absent in *mC. bovis* and present in *pC. bovis*. **(E)** Growth of *S. aureus*, expressed as area under the growth curve (AUC), in medium conditioned by *mC. bovis* or *pC. bovis* in the presence or absence of exogenous xanthine (2mM). Controls comprise unconditioned medium with *S. aureus*, conditioned medium without *S. aureus*, and medium alone. In the absence of xanthine the two conditioned media did not differ; in its presence, *mC. bovis*-conditioned medium supported significantly less *S. aureus* growth than *pC. bovis*-conditioned medium. Each dot represents an independent experiment ([N] per condition); bars represent mean ± SEM. p-values determined by two-way ANOVA.

Whole-genome alignment using the BV-BRC Genome Comparison Viewer revealed a high degree of similarity across the shared genomic backbone of the two strains (Figure 3B). A localized structural rearrangement was nonetheless identified, in which a genomic segment (2,600,000–2,620,000 bp) was translocated relative to its position in *mC. bovis*, with the corresponding flanking block repositioned further downstream in *pC. bovis*.

Bidirectional BLAST-based proteome comparison, visualized as a circular identity plot (Figure 3C), showed that most predicted proteins shared high sequence identity (>95–99.9%) between the two strains, consistent with their close phylogenetic relationship. A small subset of loci displayed reduced identity (70–90%), indicating localized strain-specific divergence or loss of a small number of genes.

Comparative pathway analysis of purine metabolism-associated enzymes similarly revealed that gene copy number was largely conserved between the two strains (Figure 3D) (Supplementary Table 1). Notably, xanthine phosphoribosyltransferase was absent in *mC. bovis* but present in *pC. bovis*, representing the only enzyme within this pathway showing a presence/absence difference between the two genomes. This finding suggests that *pC. bovis* retains an additional route for purine salvage via xanthine that is not encoded in the *mC. bovis* genome, potentially reflecting a differential metabolic capability between the two strains.

Xanthine is a purine detectable in human tears and accumulates in the tears of patients with bacterial keratitis alongside depletion of adenine and adenosine (14). We next asked if purine salvaging from xanthine may be a microbial mechanism that can influence the outcome of microbial keratitis. Because *S. aureus* is a microbe that is relevant to bacterial keratitis and relies, in part, on purine salvage to meet its nucleotide demand, we asked whether the difference in xanthine phosphoribosyltransferase between the two *Corynebacterium* strains may influence the growth of *S. aureus* in culture. We grew both strains to log phase with or without 2mM xanthine, normalized each culture to OD600 0.5, removed bacteria by filtration, and measured *S. aureus* growth in the resulting conditioned media over 36 hours as area under the curve (AUC). With xanthine, *S. aureus* reached a greater density in medium conditioned by *pC. bovis* than in medium conditioned by *mC. bovis* (*P=*0.0018; Figure 3E). Without xanthine the two conditioned media were indistinguishable, and the effect was specific to *mC. bovis*, which supported less growth once xanthine was present (*P =* 0.0071), while *pC. bovis*-conditioned medium was unchanged and xanthine alone did not alter unconditioned medium (*P =* 0.9981). Growth in *pC. bovis*-conditioned medium never exceeded unconditioned medium (*P*>0.9999). The strain difference therefore reflects xanthine-dependent restriction of *S. aureus* by *mC. bovis*, not enhancement by *pC. bovis*.

Together, *mC. bovis* and *pC. bovis* are highly similar across the genome and proteome, differing principally in a discrete structural rearrangement within a ∼50–70 kb region and in the presence or absence of xanthine phosphoribosyltransferase.

### Human *C. bovis* activates human immunity

Having established these genomic differences, we next characterized the immune responses the two isolates elicit in human cells. We reasoned that a long-term colonizer would stimulate memory immune responses, so memory responses directed against ocular bacteria would argue that these bacteria are residents rather than transient passengers. We, therefore, incubated freshly isolated PBMCs with heat-killed *mC. bovis*, *pC. bovis,* or murine *C. mastitidis* (AS1) for 72 hours, using heat-killed preparations so that differing growth rates over the culture could not confound the comparison. We then assayed the cells by flow cytometry and the supernatants by ELISA. To assess T cell activation, we monitored CD69, which marks recent antigenic stimulation (23, 24). CD69 was unchanged on CD4^+^ and CD8^+^ T cell under every condition but was significantly upregulated in the γδ T cell compartment by either *C. bovis* strains relative to media control. AS1 trended the same way without reaching significance (Figure 4A). Because we and others have found that γδ T cells respond to Corynebacteria by secreting IL-17, we also assayed the supernatants, and both *C. bovis* strains and *C. mastitidis* induced significant IL-17 (Figure 4B). Humans therefore harbor γδ T cells that recognize and respond to the eye-colonizing *C. bovis*.

**Figure 4.**
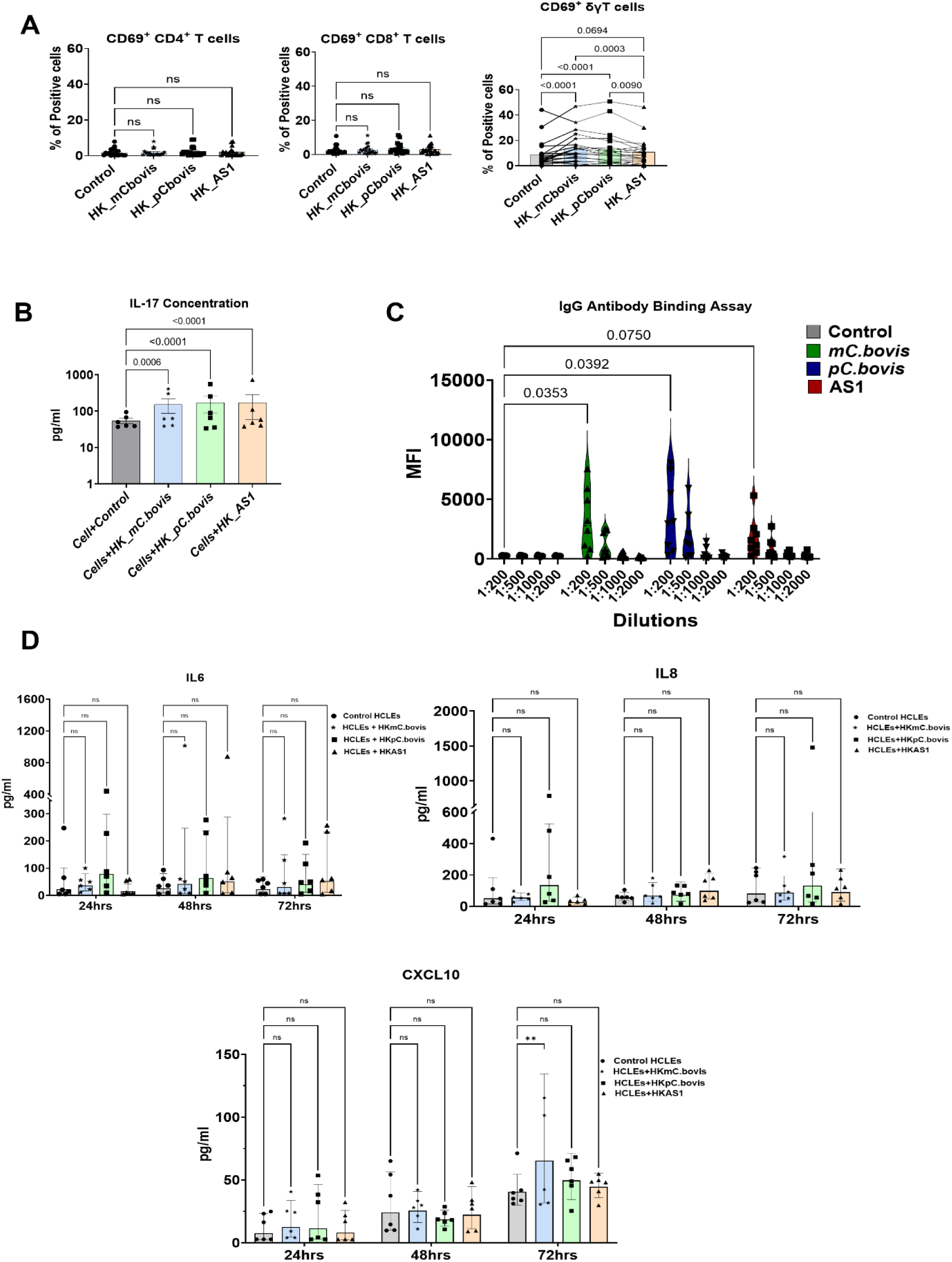
Human immune cell activation, antibody responses, and corneal epithelial cytokine production induced by heat-killed *C. bovis* strains. **(A)** Frequency of T cell activation marker (CD69⁺ CD4⁺ T cells, CD69⁺ CD8⁺ T cells, and CD69⁺ γδ T cells) were assessed following stimulation of human PBMCs with heat-killed (HK) AS1, *mC. bovis*, or *pC. bovis*, compared to unstimulated control. γδ T cell activation was significantly increased by HK_mC. bovis (p = <0.001), and HK_pC. bovis (p < 0.0001) relative to control. In Contrast neither CD4⁺ nor CD8⁺ T cell activation was significantly altered by any stimulus. Each dot represents an individual donor; bars represent mean ± SEM. **(B)** IL-17 concentration (pg/ml) in culture supernatants following stimulation of human PBMCs with heat-killed AS1, *mC. bovis*, or *pC. bovis*, compared to unstimulated control. All three heat-killed bacterial stimuli significantly increased IL-17 production relative to control (p < 0.0001 for *pC. bovis*; p = <0.001 for *mC. bovis*). Each dot represents an individual donor; bars represent mean ± SEM. **(C)** IgG antibody binding assay measuring Mean Fluorescence Intensity (MFI) of serum antibody binding to *mC. bovis*, *pC. bovis*, control, and AS1, across serial serum dilutions (1:200 to 1:2000). Significant increases in antibody binding at the 1:200 dilution were observed for *pC. bovis* (p = <0.05) and *mC. bovis* (p = <0.05) relative to control. Each dot represents an individual plasma sample. **(D)** Cytokine secretion (IL-6, IL-8, and CXCL10; pg/ml) by human corneal epithelial cells (HCLEs) following stimulation with heat-killed *mC. bovis*, or *pC. bovis*, and AS1 measured at 24, 48, and 72 hours post-stimulation, relative to unstimulated control HCLEs. IL-6 and IL-8 secretion did not significantly differ between any stimulation group and control at any time point. CXCL10 secretion was significantly increased in HCLEs stimulated with HK_mC. bovis at 72 hours relative to control (p < 0.01). Each dot represents an individual replicate; bars represent mean ± SEM.

B cells, like T cells, require antigen-specific activation before they secrete antibody, so we next assayed circulating bacterium-specific antibodies. We incubated diluted serum from healthy ocular subjects with each strain, stained with fluorescent anti-human IgG, and detected antibody-bound bacteria by flow cytometry. Serum bound both *C. bovis* strains significantly (*mC. bovis* MFI 3271 ± 2316 a.u.; *pC. bovis* MFI 3660 ± 2681 a.u.) relative to control (MFI 275 ± 29.1 a.u.), while murine C. mastitidis trended the same way without reaching significance (MFI 1,803 ± 1,424 a.u.; Figure 4C). Taken with the γδ T cell data, healthy ocular subjects mount adaptive immune responses to *C. bovis*, which again points to long-term exposure.

Other cells likely respond to ocular commensals in vivo, so we asked how the corneal epithelium reacts to these strains. Human corneal epithelial (HCLE) cells were incubated with heat-killed *mC. bovis*, *pC. bovis*, or AS1 for 24, 48, and 72 hours, and we measured IL-6, IL-8, and CXCL10 in the supernatants by ELISA against unstimulated controls, choosing these three for their central roles in epithelial signaling and inflammatory recruitment. Neither IL-6 nor IL-8 changed with any strain at any time point (Figure 4D), which suggests these commensals do not trigger classical pro-inflammatory epithelial signaling, at least under the conditions examined here. By 72 hours, however, *mC. bovis* significantly raised CXCL10 (80 ± 49 pg/mL vs. 42 ± 13.5 pg/mL in controls), while neither *pC. bovis* nor AS1 elicited a CXCL10 response at any time point (Figure 4D). These strains are therefore not excessively inflammatory, supporting the idea that colonization would not drive pathogenic inflammation in human hosts.

### Human conjunctiva harbors immune cells that may recognize ocular surface bacteria

In mice, we have previously shown that immune cells traffic to the conjunctiva in response to the status of the ocular microbiome (15). Knowing the dynamic nature of the conjunctiva in response to stimulation (7, 25, 26), we sought to understand the identities of cells present in the human conjunctiva with healthy ocular surface, so we isolated conjunctival cells from subjects at the Miami and Pittsburgh sites non-invasively using an EyePrim device (Opia Technologies), and comprehensively analyzed immune cell populations using flow cytometry.

Analysis of CD45□ total immune cells revealed a diverse immune landscape at the ocular surface (Figure 5A). αβ T cells comprised the dominant lymphocyte population, with both CD4□ helper and CD8□ cytotoxic subsets well represented across all samples. CD8□ T cells were more abundant in the conjunctiva compared to CD4+ T cells. These results agree with previous results and support the notion that the eye is routinely challenged by viruses from the environment (27–29). CD8^+^ T cells were significantly more frequent in Miami samples than in Pittsburgh samples (Figure 5B). This may reflect the heightened antigen exposure of Miami’s subtropical climate, where allergen and microbial burden are known to raise ocular surface immune activation.

**Figure 5.**
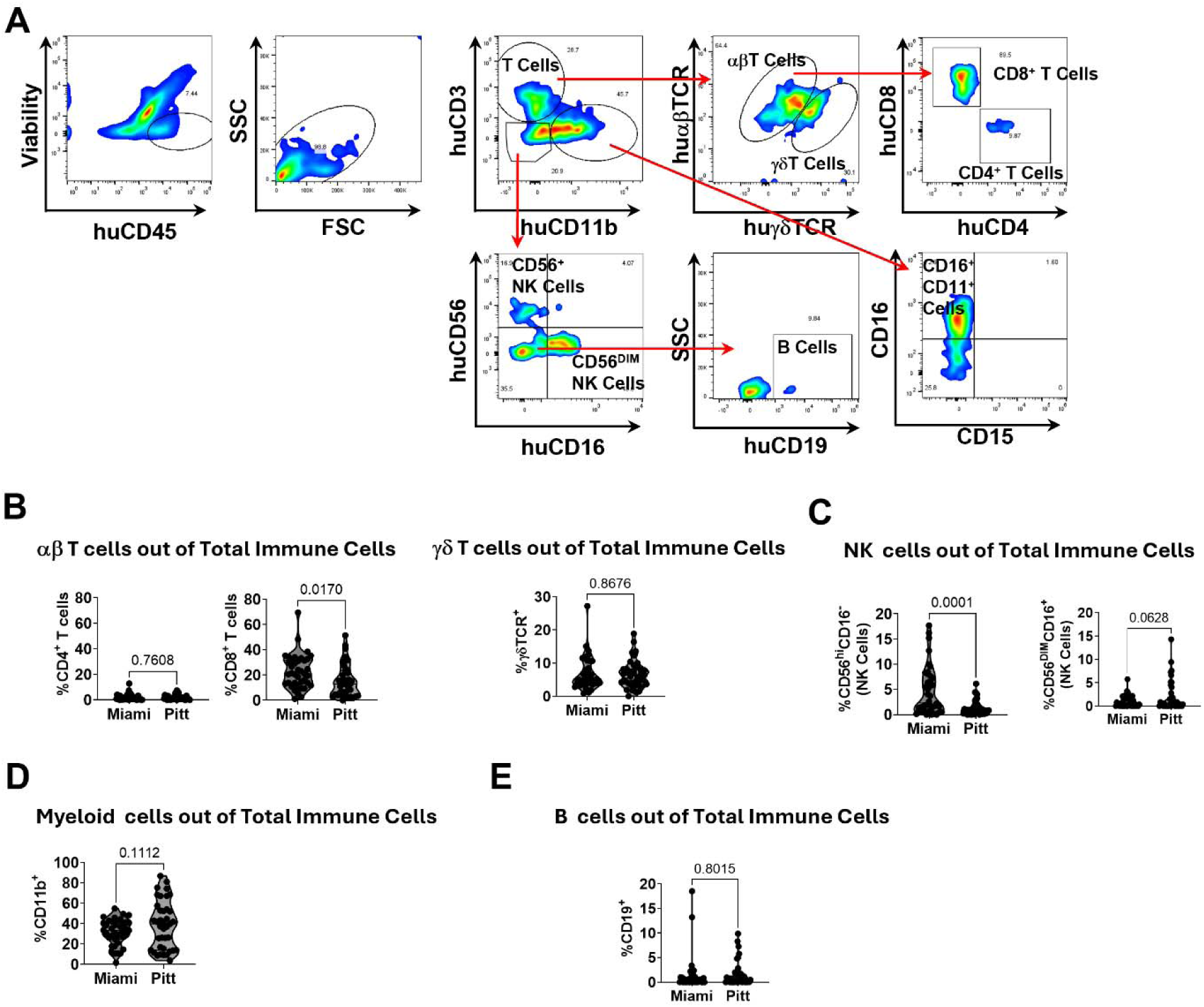
Human conjunctiva harbors immune cells that may recognize ocular surface bacteria. **(A)** Representative flow cytometric gating strategy used to identify immune cell populations in human conjunctival samples. Within the huCD45⁺ population, T cells (huCD3⁺huCD11b⁻) were further subdivided into αβ T cells and γδ T cells, with αβ T cells further classified into CD4⁺ and CD8⁺ T cell subsets. The huCD3⁻huCD11b⁺ population was analyzed for NK cells (CD56^+^), B cells (huCD19⁺), and CD16⁺CD11⁺ myeloid cells based on CD16 and CD15 expression. **(B-E)** Frequency of αβ T cells (%CD4⁺ and %CD8⁺ T cells), γδ T cells, NK cell subsets (%CD56^hi^CD16⁻ and %CD56^dim^CD16⁺), myeloid cells (%CD11b⁺), and B cells (%CD19⁺) expressed as a percentage of total immune cells, in conjunctival samples from Miami and Pittsburgh sites. Each dot represents an individual sample; p-values determined by Welch’s t-test.

Consistent with the *in vitro* data, resident γδ T cells were detected in healthy conjunctival samples from both sites, reinforcing their identity as components of the ocular mucosal immune system. γδT cell frequency did not differ between the Miami and Pittsburgh cohorts (Figure 5C), suggesting that the γδ T cell population is maintained as a stable population of immune cells regardless of geographic or environmental context. Natural killer (NK) cells were present in healthy conjunctiva at both sites. The immunoregulatory CD56^hi^CD16 subset differed significantly between sites, while cytotoxic CD56^dim^CD16 NK cells only trended toward a difference (Figure 5C). This divergence points to site-specific tuning of immunoregulatory innate responses, likely driven by the distinct environmental and microbial landscapes each population encounters. Myeloid and B cell frequencies, by contrast, were comparable between the two cohorts (Figures 5D–E).

### Human Corynebacteria colonize the mouse eye and stimulate protective immunity

To test whether the mouse eye can model host interactions with human ocular microbes, we colonized mice with human-derived *Corynebacterium bovis* and monitored them over three weeks (Figure 6A). At the endpoint, we quantified bacterial burden and profiled ocular immune cells by flow cytometry.

**Figure 6:**
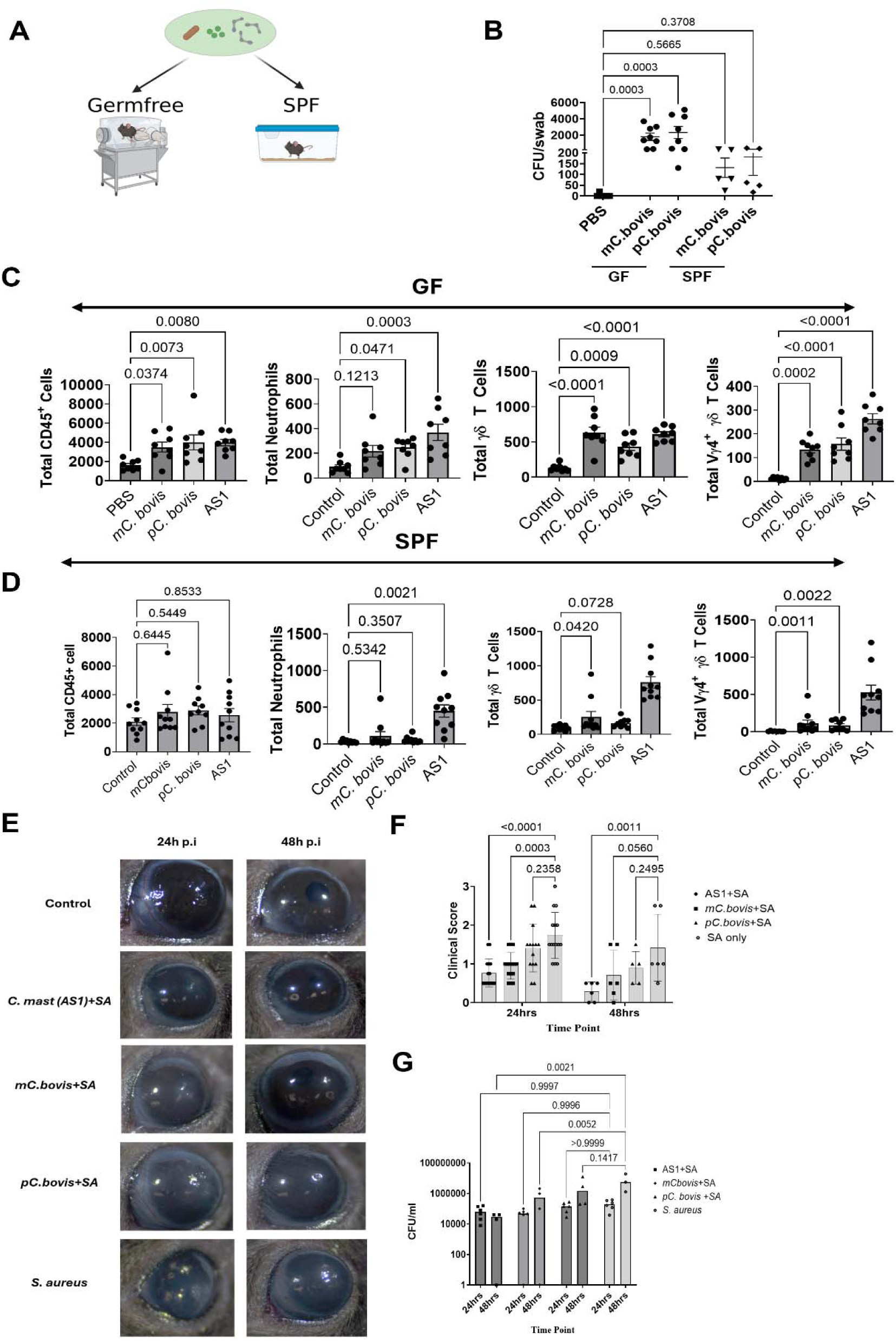
Ocular colonization with *C. bovis* strains induces γδ T cell responses in germ-free mice and confers protection against *S. aureus* keratitis. (A) Schematic of experimental colonization strategy: Germ-free (GF) and Specific-pathogen-free (SPF) mice were topically inoculated on the ocular surface with *mC. bovis,* or *pC. bovis*. (B) Bacterial burden (CFU/swab) recovered from the ocular surface of GF (n= 8/group) and SPF mice (n=5/group) following inoculation with *mC. bovis*, *pC. bovis,* PBS (control). Each dot represents an individual mouse; bars represent mean ± SEM. p-values determined by Kruskal-Wallis test used with multiple comparisons. (C) Immune cells (Total CD45⁺, neutrophils, γδ T, and Vγ4⁺ γδ T cells) quantified at the ocular surface of GF mice following colonization with *mC. bovis*, *pC. bovis*, PBS (control), and AS1 (*C. mastitidis*, positive control). Each dot represents an individual mouse (n= 8/group); bars represent mean ± SEM. p-values determined by one-way ANOVA test. (D) Immune cells quantified at the ocular surface of SPF mice following colonization with *mC. bovis*, *pC. bovis*, control, and AS1. Each dot represents an individual mouse (n=10/group); bars represent mean ± SEM. p-values determined by one-way ANOVA test. (E) Representative slit-lamp images of the ocular surface at 24 and 48 hours post-infection (p.i.) with *S. aureus* (SA) in mice pre-colonized with *mC. bovis*, *pC. bovis*, control (PBS), AS1, and infected with *S. aureus* (SA) alone. (F) Clinical scores of *S. aureus* keratitis severity at 24 and 48 hours p.i. in mice pre-colonized with *mC. bovis* (24h n=15; 48h n= 6)*, pC. bovis* (24h n=15; 48h n= 6), AS1 (24h n=15; 48h n= 6), prior to SA challenge, compared to SA-only infected controls (24h n=15; 48h n= 6). Each dot represents an individual mouse; bars represent mean ± SEM. p-values determined by Two-way ANOVA test. (G) Corneal bacterial burden (CFU/ml) at 24 and 48 hours post-*S. aureus* infection in mice pre-colonized with *mC. bovis* (24h n=6; 48h n= 4)*, pC. bovis* (24h n=6; 48h n= 4), and AS1 (24h n=6; 48h n= 4) compared to SA-only infected controls (24h n=6; 48h n= 4). Each dot represents an individual mouse; bars represent mean ± SEM. p-values determined by Two-way ANOVA test.

Germ-free (GF) and specific pathogen-free (SPF) mice were inoculated with *mC. bovis* (Miami) or *pC. bovis* (Pittsburgh), with PBS as a control. Both strains colonized the ocular surface, but far more efficiently in GF than in SPF mice. In GF animals, *mC. bovis* reached a mean of 1.80×10³ CFU and *pC. bovis* 2.33×10³ CFU, which were significantly higher than the PBS control (Figure 6B). SPF mice were also colonized, though at lower burden and with no difference between strains (*mC. bovis* 1.31×10² CFU; *pC. bovis* 1.80×10² CFU). We concluded from these data that the resident ocular microbiota limits, but does not prevent, the establishment of human *C. bovis* colonization at the ocular surface.

We next asked whether colonization with *C. bovis* stimulated local immunity compared to PBS and the established murine commensal, *C. mastitidis* (served as a positive control). In GF mice, both *mC. bovis* and *pC. bovis* drove the recruitment of γδ T cells and the Vγ4 subset (Figure 6C) to the ocular surface. Given the role of Vγ4 cells in IL-17 production and epithelial defense, this points to a targeted, commensal-driven response. Neutrophil recruitment, by contrast, was unchanged for either strain compared to the PBS control suggesting that unlike *C. mastitidis*, *C. bovis* primarily drives lymphoid over a myeloid responses.

In SPF mice, these responses were attenuated, but not abolished. Notably, *mC. Bovis* significantly expanded both total γδ and Vγ4□γδ T cells in SPF animals, whereas *pC. Bovis* altered neither population, and neither strain changed total CD45□ cell or neutrophil numbers (Figure 6D). Under conventional microbial conditions the γδ T cell response to *C. bovis* is therefore strain restricted, with only *mC. bovis* engaging the Vγ4□ compartment.

We then asked whether this modest response was protective. *Staphylococcus aureus* is a leading cause of bacterial conjunctivitis and keratitis, so we tested whether prior commensal colonization guards against it. SPF C57BL/6 mice colonized with *mC. bovis*, *pC. bovis*, or AS1 for three weeks were challenged with 1×10□ CFU *S. aureus*. Corneal opacity appeared within 24 hours in all infected mice, while uninfected eyes stayed clear and SA-only eyes developed pronounced opacity (Figure 6E). *mC. bovis* and *C. mastitidis* both correlated with reduced opacity at 24 hours. For *C. mastitidis*, this significantly persisted for 48 hours, but for *mC. bovis* there was a trend towards lowered opacity (*P* = 0.0560; Figures 6E–F). *pC. bovis* trended in a similar way; however, there was no significance at any time point even though *pC. bovis* treated groups harbored expanded γδ and Vγ4□ populations in germ-free mice as efficiently as *mC. bovis* (Figure 6C).

Bacterial burden followed a similar course to the clinical opacity. CFU rose from 24 to 48 hours across all groups, consistent with active *S. aureus* replication, but by 48 hours colonized and uncolonized eyes diverged. *C. mastitidis*- and *mC. bovis*–colonized eyes harbored significantly less *S. aureus* compared to *pC. bovis* and uncolonized controls (Figure 6G). Together, these data show that *Corynebacterium*-mediated protection is strain dependent, which raises the question for future studies of which *C. bovis* factors are required to confer protection from pathogenic bacterial infection.

## DISCUSSION

Whether the ocular surface sustains a stable, functional microbial community or is merely seeded transiently from adjacent skin and the environment has remained unresolved, largely because the paucibacterial nature of the site leaves single-method surveys vulnerable to reagent contamination and sampling errors. By pairing culture-independent sequencing with culturomics across two geographically distinct cohorts, we show that the healthy human conjunctiva harbors a reproducible core microbiome that is dominated by *Staphylococcus* and *Corynebacterium* and is relatively consistent at a genus level across individuals and locations. We also found that this community is actively engaged by the host rather than passively tolerated, and that a previously defined protective network of commensal/host interactions driven by γδ T cell IL-17 may be recapitulated in the human host (15). Together, our data move the ocular microbiome from correlative assessment of detectable taxa toward a defined, immunologically integrated, and functionally consequential feature of ocular surface microbe/host homeostasis.

The credibility of defining any low-biomass “core” depends on the methods used to define it, and prior disagreements in the field, including longitudinal work that recovered no conserved taxa across healthy individuals, fairly reflect the limits of any single approach. Since most prior efforts to define an ocular core have relied on DNA, sequencing results will detect DNA from both viable and non-viable organisms on the surface and within reagents as contaminants, while culture-based methods underestimate diversity (20, 30). Here, we are analyzing microbiota based on16S rRNA sequences (as opposed to 16S rRNA gene sequences from DNA). Therefore, we are examining taxa that are transcribing or maintaining stable rRNA, thus weighing identification of core taxa towards those taxa that are metabolically active(31–33). Coupling these culture independent and culture dependent methods together provided the orthogonal validation needed to identify Staphylococcus and Corynebacterium as the dominant members of the core ocular surface microbiome. Notably, while genus-level composition was conserved between Pittsburgh and Miami, consistent with 16S surveys spanning tropical and continental climates (1, 34–38) that routinely detect *Corynebacterium, Staphylococcus*, and *Propionibacterium*, species and strain level differences emerged only when culturomics was layered onto sequencing. Taken together, geography does not change which genera can occupy the niche, but it does shape which species or strains of a genus such as Corynebacterium do so.

This distinction points to a broader principle: a genus-anchored functional core may maintain the local resident immune landscape. Indeed, we found that geography influenced significant differences in CD8+ T cell and NK cell frequencies within the conjunctiva; however, γδ T cells, a population we and others have previously shown recognize and respond to Corynebacteria, are maintained at similar frequencies in both cohorts (39, 40). The core microbiome and its primary immune correlate therefore co-exist at the ocular surface in both communities.

The conserved nature between human and mouse host-microbiome relationships is important because it provides a model for studying ocular host-microbiome interactions in functional depth rather than by correlation. Human γδ T cells, but not CD4^+^ or CD8^+^ T cells, upregulated CD69 in response to both murine *C. mastitidis* and the two human *C. bovis* strains, which argues for a microbe-specific rather than a general antibacterial response, consistent with previous findings. Release of IL-17 from immune cells into *C. bovis* stimulated cultures and detectable levels of circulating *C. bovis*-specific IgG in healthy subject sera points towards a holistic memory immune response towards this microbe. We reasoned that, together, these observations of long-term immunity suggest that *C. bovis* is a human colonizing microbe.

At the epithelium, the absence of inflammatory cytokines like IL-6 and IL-8 indicates that these commensals do not trigger classical inflammatory signaling (41, 42). The modest induction of CXCL10 could be a recruitment factor that brings γδ T cells to the tissue *in vivo*, which would link epithelial sensing to the establishment of resident lymphocytes.

Functionally, prior colonization with human *C. bovis* reduced both the clinical severity and bacterial burden of *S. aureus* keratitis, with *mC. bovis* driving an approximately ten-fold reduction in corneal CFU. This extends our foundational work on *C. mastitidis* by extending the protective benefits of ocular colonization of Corynebacteria to *S. aureus*, a leading Gram-positive cause of sight-threatening keratitis not previously tested and to a human-derived commensal. Our working model currently parallels other reports of *Corynebacterium*-mediated antagonism of *S. aureus* in the airway, suggesting a conserved cross-barrier defense strategy (43–47).

The divergence between the two human isolates is instructive. In germ-free mice, *pC. bovis* colonized and expanded γδ and Vγ4□ populations at least as efficiently as *mC. bovis*, so protection cannot be read from colonization efficiency or from bulk γδ expansion under gnotobiotic conditions. The distinction emerges under conventional conditions. In SPF mice, the setting in which we assessed *S. aureus* challenge, only *mC. bovis* recruited γδ and Vγ4 compartments to the ocular surface. Notably, *mC. bovis* was also the only strain to induce epithelial-derived CXCL10, a CXCR3 ligand positioned to support recruitment of γδ T cells into the tissue (39, 40). Protective immune-centric mechanisms, therefore, may be dependent on the ability of ocular microbiome components recruit γδ T cells to the ocular surface and at other potential epithelial barriers (15, 41, 42). Because an intact microbiota blunts commensal-driven ocular immunity overall, the capacity to engage this compartment despite that competition appears to be a strain-level property rather than a general feature of ocular *Corynebacteria*.

A second, non-immune axis may also contribute to ability of the ocular surface to resist infection. A major genomic difference we detected between the two strains centered around xanthine phosphoribosyltransferase, which is absent in *mC. bovis* and present in *pC. bovis*. The functional consequence of this is that, in the presence of xanthine, *mC. bovis* supported less *S. aureus* growth than *pC. bovis*. Because xanthine is detectable in human tears (14), and that *S. aureus* depends, in part, on purine salvage, a strain-specific effect on the local purine pool offers a plausible route by which commensal *Corynebacteria* may modify pathogen fitness independently of the host response. A possible explanation for this is that a commensal that cannot metabolize xanthine would draw more heavily on other purine sources and create a less permissive environment behind for pathogenic bacteria like *S. aureus*. The modest magnitude of the *in vitro* effect suggests that this metabolic difference is a candidate modifier of commensal-mediated protection rather than an explanation, and a combination of both the immune and non-immune factors is required for maximal ocular surface protection.

Limitations in the study include: **1)** The human immune data linking γδ activation and circulating IgG are correlative, but do not themselves demonstrate human mechanisms. **2)** The colonization difference between germ-free and SPF mice also warrants further work, since it remains unclear why bacteria colonize the germ-free eye more efficiently. Germ-free animals lack both competition from an endogenous microbiome and a fully educated immune system, but we did not test which accounts for the difference. **3)** Finally, our use of heat-killed bacteria, a cross-sectional design that infers rather than longitudinally tracks stability, and recruitment from two U.S. sites all constrain generalizability.

Collectively, these findings reframe the ocular microbiome not as a passive community to be tolerated but as an active participant in mucosal immune homeostasis, and they identify human-derived commensal *Corynebacterium* as a rational, mechanism-based tool for protecting the ocular surface. Given the rising burden of antimicrobial resistance and the severe morbidity of bacterial keratitis, a commensal-anchored γδ–IL-17 axis offers a plausible foundation for microbiome-guided prophylaxis in high-risk settings such as contact lens wear, ocular surgery, and trauma, a prospect that motivates causal dissection of the axis, definition of the strain determinants of protection, and longitudinal validation in humans.

## Author contributions

AJS, VJ, and AG conceptualized and designed the study. JG, BD, SMM, VJ, AG, AW, and JB Human Sample Collection; BM, KL, and AF Sequencing, data analysis and data interpretation SKR, and MM Culturomics; JG Invitro experiments; JG and AJS Mice experiments; JG, BD, MM, BM, KL, AF and AJS analyzed the data. JG and BD prepared the initial draft of the manuscript. AG, AJS, BD, JG, SMM, and VJ wrote the manuscript, and all authors reviewed and editing the final text.

## References

1. Doan T, Akileswaran L, Andersen D, Johnson B, Ko N, Shrestha A, et al. Paucibacterial Microbiome and Resident DNA Virome of the Healthy Conjunctiva. Invest Ophthalmol Vis Sci. 2016;57(13):5116–26.

2. Lai J, Rigas Y, Kantor N, Cohen N, Tomlinson A, St Leger AJ, et al. Living with your biome: how the bacterial microbiome impacts ocular surface health and disease. Expert Rev Ophthalmol. 2024;19(2):89–103.

3. Zilliox MJ, Gange WS, Kuffel G, Mores CR, Joyce C, de Bustros P, et al. Assessing the ocular surface microbiome in severe ocular surface diseases. Ocul Surf. 2020;18(4):706–12.

4. Herzog EL, Kreuzer M, Zinkernagel MS, and Zysset-Burri DC. Challenges and insights in the exploration of the low abundance human ocular surface microbiome. Front Cell Infect Microbiol. 2023;13:1232147.

5. Niederkorn JY, and Kaplan HJ. Rationale for immune response and the eye. Chem Immunol Allergy. 2007;92:1–3.

6. Niederkorn JY. Cornea: Window to Ocular Immunology. Curr Immunol Rev. 2011;7(3):328–35.

7. Steven P, Schwab S, Kiesewetter A, Saban DR, Stern ME, and Gehlsen U. Disease-Specific Expression of Conjunctiva Associated Lymphoid Tissue (CALT) in Mouse Models of Dry Eye Disease and Ocular Allergy. Int J Mol Sci. 2020;21(20).

8. Dunlap DG, Yang L, Qin S, Li K, Fitch A, Huang L, et al. Magnetic-activated cell sorting identifies a unique lung microbiome community. Microbiome. 2023;11(1):117.

9. Kitsios GD, Li K, Blacka S, Fitch A, Jacobs J, Naqvi A, et al. Oral and gut microbiota relate to symptom subphenotypes in long COVID, independent of viral persistence. iScience. 2025;28(11):113628.

10. Li K, Bihan M, and Methe BA. Analyses of the stability and core taxonomic memberships of the human microbiome. PLoS One. 2013;8(5):e63139.

11. Phelps CM, Shapira JH, Laughlin CR, and Meisel M. Detection of viable commensal bacteria in murine melanoma tumors by culturomics. STAR Protoc. 2023;4(3):102492.

12. Bose T, Hou A, Lee R, Tong L, and Chandy KG. A Non-invasive Way to Isolate and Phenotype Cells from the Conjunctiva. J Vis Exp. 2017(125).

13. Wattam AR, Davis JJ, Assaf R, Boisvert S, Brettin T, Bun C, et al. Improvements to PATRIC, the all-bacterial Bioinformatics Database and Analysis Resource Center. Nucleic Acids Res. 2017;45(D1):D535–D42.

14. Shrestha GS, Vijay AK, Stapleton F, White A, Pickford R, and Carnt N. Human tear metabolites associated with nucleoside-signalling pathways in bacterial keratitis. Exp Eye Res. 2023;228:109409.

15. St Leger AJ, Desai JV, Drummond RA, Kugadas A, Almaghrabi F, Silver P, et al. An Ocular Commensal Protects against Corneal Infection by Driving an Interleukin-17 Response from Mucosal gammadelta T Cells. Immunity. 2017;47(1):148–58 e5.

16. Rigas Y, Treat BR, Shane J, Shanks RMQ, and St Leger AJ. Genetic Manipulation of Corynebacterium mastitidis to Better Understand the Ocular Microbiome. Invest Ophthalmol Vis Sci. 2023;64(2):19.

17. Rizzuto V, Settino M, Stroffolini G, Covello G, Vanags J, Naccarato M, et al. Ocular surface microbiome: Influences of physiological, environmental, and lifestyle factors. Comput Biol Med. 2025;190:110046.

18. Apuy Rodriguez F, Chacon Quiros M, Alvarado Fernandez ML, Alvarado Mora ML, and Vanegas Navarro P. Microbiome of the Ocular Surface: Resident or Transient Ecosystem? Cureus. 2025;17(8):e89487.

19. Ahn J, and Hayes RB. Environmental Influences on the Human Microbiome and Implications for Noncommunicable Disease. Annu Rev Public Health. 2021;42:277–92.

20. Salter SJ, Cox MJ, Turek EM, Calus ST, Cookson WO, Moffatt MF, et al. Reagent and laboratory contamination can critically impact sequence-based microbiome analyses. BMC Biol. 2014;12:87.

21. Glassing A, Dowd SE, Galandiuk S, Davis B, and Chiodini RJ. Inherent bacterial DNA contamination of extraction and sequencing reagents may affect interpretation of microbiota in low bacterial biomass samples. Gut Pathog. 2016;8:24.

22. Duan T, Rosenbaum A, Chandra V, Tiraboschi L, Rescigno M, McAllister F, et al. Toward a consensus on the tumor microbiota: Evidence, standards, and interpretation. Cancer Cell. 2026;44(4):696–702.

23. Craston R, Koh M, Mc Dermott A, Ray N, Prentice HG, and Lowdell MW. Temporal dynamics of CD69 expression on lymphoid cells. J Immunol Methods. 1997;209(1):37–45.

24. Cibrian D, and Sanchez-Madrid F. CD69: from activation marker to metabolic gatekeeper. Eur J Immunol. 2017;47(6):946–53.

25. Knop E, and Knop N. The role of eye-associated lymphoid tissue in corneal immune protection. J Anat. 2005;206(3):271–85.

26. Liu Y, Zhu R, Jin X, Wang Y, Shi Y, Zhang N, et al. Activation of Conjunctiva-Associated Lymphoid Tissue in Patients With Infectious Keratitis Using In Vivo Confocal Microscopy. Invest Ophthalmol Vis Sci. 2021;62(10):27.

27. Loi JK, Alexandre YO, Senthil K, Schienstock D, Sandford S, Devi S, et al. Corneal tissue-resident memory T cells form a unique immune compartment at the ocular surface. Cell Rep. 2022;39(8):110852.

28. Kuklinski EJ, Yu Y, Ying GS, Asbell PA, and Group DSR. Association of Ocular Surface Immune Cells With Dry Eye Signs and Symptoms in the Dry Eye Assessment and Management (DREAM) Study. Invest Ophthalmol Vis Sci. 2023;64(12):7.

29. Downie LE, Zhang X, Wu M, Karunaratne S, Loi JK, Senthil K, et al. Redefining the human corneal immune compartment using dynamic intravital imaging. Proc Natl Acad Sci U S A. 2023;120(31):e2217795120.

30. Emerson JB, Adams RI, Roman CMB, Brooks B, Coil DA, Dahlhausen K, et al. Schrodinger’s microbes: Tools for distinguishing the living from the dead in microbial ecosystems. Microbiome. 2017;5(1):86.

31. Campbell BJ, Yu L, Heidelberg JF, and Kirchman DL. Activity of abundant and rare bacteria in a coastal ocean. Proc Natl Acad Sci U S A. 2011;108(31):12776–81.

32. Li R, Tun HM, Jahan M, Zhang Z, Kumar A, Dilantha Fernando WG, et al. Comparison of DNA-, PMA-, and RNA-based 16S rRNA Illumina sequencing for detection of live bacteria in water. Sci Rep. 2017;7(1):5752.

33. Li R, Tun HM, Jahan M, Zhang Z, Kumar A, Dilantha Fernando WG, et al. Author Correction: Comparison of DNA-, PMA-, and RNA-based 16S rRNA Illumina sequencing for detection of live bacteria in water. Sci Rep. 2018;8(1):17427.

34. Wen X, Miao L, Deng Y, Bible PW, Hu X, Zou Y, et al. The Influence of Age and Sex on Ocular Surface Microbiota in Healthy Adults. Invest Ophthalmol Vis Sci. 2017;58(14):6030–7.

35. Kim JW, Lee JS, Kim JH, Jeong JW, Lee DH, and Nam S. Comparison of Microbiota Variation in Korean Healthy Adolescents with Adults Suggests Notable Maturity Differences. OMICS. 2018;22(12):770–8.

36. Graham JE, Moore JE, Jiru X, Moore JE, Goodall EA, Dooley JS, et al. Ocular pathogen or commensal: a PCR-based study of surface bacterial flora in normal and dry eyes. Invest Ophthalmol Vis Sci. 2007;48(12):5616–23.

37. Dong Q, Brulc JM, Iovieno A, Bates B, Garoutte A, Miller D, et al. Diversity of bacteria at healthy human conjunctiva. Invest Ophthalmol Vis Sci. 2011;52(8):5408–13.

38. Zhou Y, Holland MJ, Makalo P, Joof H, Roberts CH, Mabey DC, et al. The conjunctival microbiome in health and trachomatous disease: a case control study. Genome Med. 2014;6(11):99.

39. Yoon KC, Park CS, You IC, Choi HJ, Lee KH, Im SK, et al. Expression of CXCL9, -10, -11, and CXCR3 in the tear film and ocular surface of patients with dry eye syndrome. Invest Ophthalmol Vis Sci. 2010;51(2):643–50.

40. Carreira M, Barros L, Cruz RM, Ferreira C, Costa C, Ferreira CM, et al. CXCR3 signaling promotes Delta One T cell recruitment and antitumor efficacy in colorectal cancer. J Immunother Cancer. 2026;14(5).

41. Nielsen MM, Witherden DA, and Havran WL. gammadelta T cells in homeostasis and host defence of epithelial barrier tissues. Nat Rev Immunol. 2017;17(12):733–45.

42. Zhang H, Carnevale G, Polese B, Simard M, Thurairajah B, Khan N, et al. CD109 Restrains Activation of Cutaneous IL-17-Producing gammadelta T Cells by Commensal Microbiota. Cell Rep. 2019;29(2):391–405 e5.

43. Arrieta MC, and Finlay BB. The commensal microbiota drives immune homeostasis. Front Immunol. 2012;3:33.

44. Pickard JM, Zeng MY, Caruso R, and Nunez G. Gut microbiota: Role in pathogen colonization, immune responses, and inflammatory disease. Immunol Rev. 2017;279(1):70–89.

45. Coates M, Lee MJ, Norton D, and MacLeod AS. The Skin and Intestinal Microbiota and Their Specific Innate Immune Systems. Front Immunol. 2019;10:2950.

46. Tian K, Jing D, Lan J, Lv M, and Wang T. Commensal microbiome and gastrointestinal mucosal immunity: Harmony and conflict with our closest neighbor. Immun Inflamm Dis. 2024;12(7):e1316.

47. Tamkin E, Lorenz BP, McCarty A, Fulte S, Eisenmesser E, Horswill AR, et al. Airway Corynebacterium interfere with Streptococcus pneumoniae and Staphylococcus aureus infection and express secreted factors selectively targeting each pathogen. Infect Immun. 2025;93(2):e0044524.

