## Supplementary figures and images for "Defining the Functional Consequence of the Human Ocular Microbiome"

### Supplemental Figure 1

Supplemental Figure 1. Significant correlations of the microbiome signature with co-variates.

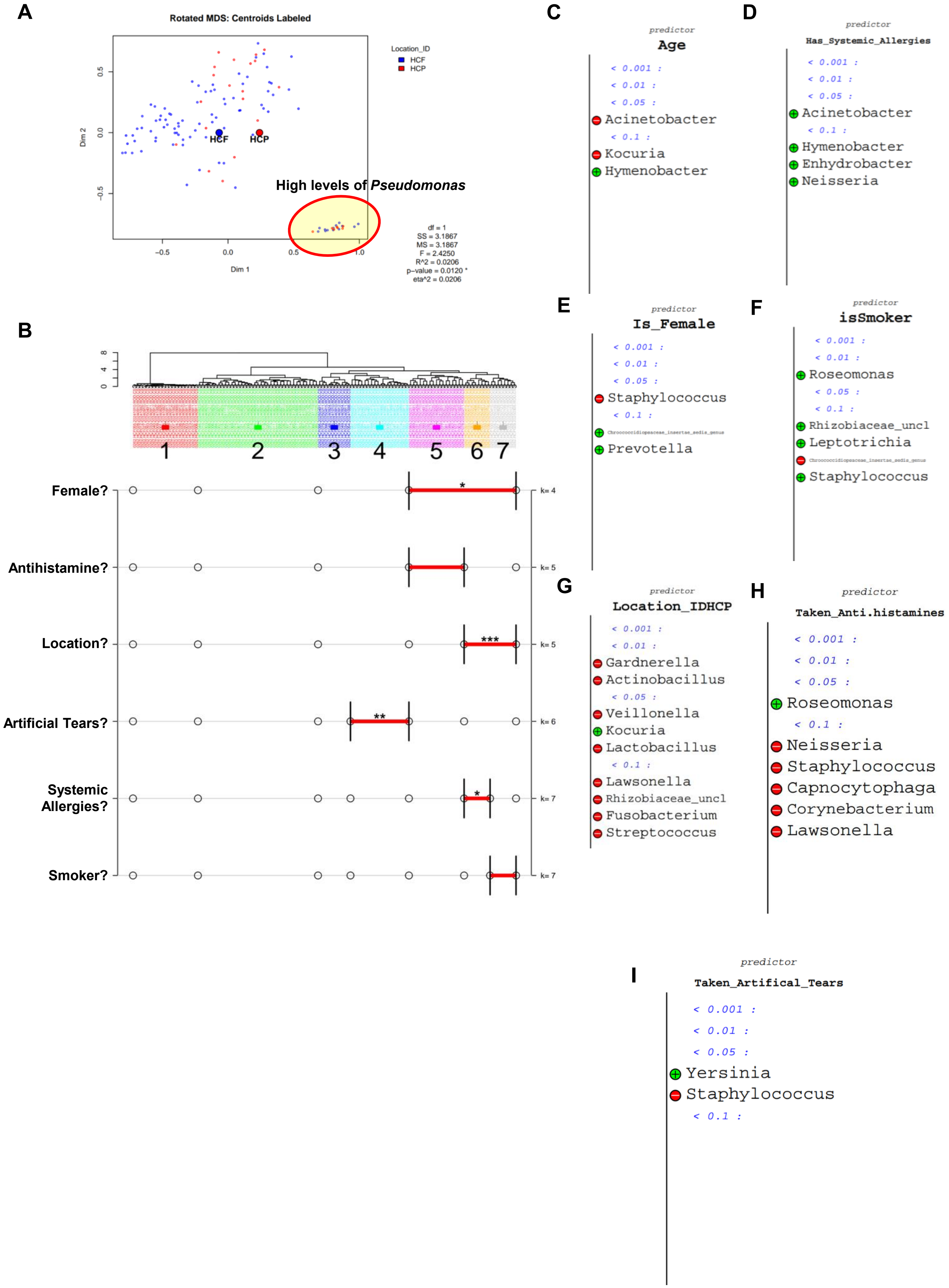
